# Canonical Correlation Analysis and Multi-Channel Cardiography Improve Artefact Cleaning in Heartbeat-Locked Analyses

**DOI:** 10.64898/2026.07.14.738519

**Authors:** Carmen Vidaurre, Maria Azanova, Ksenia Germanova, Eva Greschke, Tim Paul Steinfath, Ulrich Laufs, Tobias Uhe, Arno Villringer, Vadim V. Nikulin

## Abstract

Cardiac artefacts are problematic for neurophysiological analyses, especially for heartbeat-locked brain responses where neural activity and artefacts co-occur in time. Traditional approaches using single-channel electrocardiography for artefact removal do not fully capture the multidimensional spread of cardiac fields. Moreover, even if several channels are recorded, no established methods exists for integrating them for artefact removal. Here, we propose a multivariate approach based on Canonical Correlation for cardiac artefact removal and report its effectiveness in electroencephalography recorded simultaneously with a custom 15-channel electrocardiography in 14 participants. We quantify cleaning quality via residual R-peak artefact and neural signal preservation via alpha power. Canonical Correlation systematically outperforms the traditional Independent Component Analysis in both reducing R-peak artefacts and preserving alpha power. By analysing all possible channel combinations, we found that neck or supraclavicular electrodes improve cleaning when only a few channels are available. With four or five channels, precordial electrodes in combination with limb or supraclavicular locations provide a performance comparable to the 15-channel setup. While these findings require validation in other datasets, we outline clear decision criteria for cleaning efficiency and show that canonical correlation is a reliable approach for multi-channel cardiac artefact removal.

## 1 Introduction

There is an increasing interest in how bodily signals shape cognition and behavior [1], and cardiac activity has received particular attention [2]. On the one hand, researchers investigate the brain’s electrical activity in response to heartbeats (termed heartbeat-evoked response, HER, or potential, HEP), and relate it to cognitive functions [3, 4]. On the other hand, studies of behavioural and neural activity across different phases of the cardiac cycle demonstrate a dynamic interplay between brain, body, and behavior [5, 6]. Together, these lines of research position heartbeat-locked analysis as a central tool in neuroscience of interoception, autonomic-cortical coupling, and embodied cognition.

A fundamental challenge for this research is the contamination of electrophysiological recordings by cardiac field artefacts (CFA). The electromagnetic signal generated by the beating heart spreads nearly instantaneously through volume conduction to scalp electrodes in electroencephalography (EEG) or sensors in magnetoencephalography (MEG), producing an artefact that overlaps with neural oscillations both spectrally and temporally [3, 7–9]. On the other hand, the mechanical consequences of the heartbeat, including pulsation of vessels and ballistocardiographic effects arising from head movement induced by the cardiac pulse wave, introduce time-locked but delayed artefact components that differ in their spatiotemporal structure from the direct electrical projection [10, 11].

These artefacts are especially problematic for heartbeat-locked analyses, complicating the interpretation of whether differences between experimental conditions reflect genuine neural effects [12]. Similarly, since CFA varies systematically across the cardiac cycle with rotation and movement of the heart, the confounding influence on measured electrophysiological signals is not constant, but fluctuates in a way that can mimic or mask true cardiac-cycle modulations of neural activity. Even beyond heart-brain research, when heartbeats are coupled to external stimuli, whether intentionally or not, researchers must rule out the possibility that differences in stimulus-evoked activity stem from overlapping cardiac artefacts [13, 14]. Furthermore, there is accumulating evidence that cardiac artefact removal may critically influence the estimation of the aperiodic (1/f) component of the power spectrum [15, 16].

Various methods have been proposed for cardiac artefact removal [17–20]. The most widely used of these is independent component analysis (ICA), a blind source separation method that decomposes the EEG into statistically independent sources, from which cardiac-related components are identified and projected out of the data [12, 21]. Despite its prevalence, ICA decomposes the EEG without reference to cardiac activity. Hence, there is no guarantee that resulting components cleanly isolate artefacts from neural signals, especially because they share spatiotemporal and frequency characteristics [8, 22].

Another standard design choice is to record a single electrocardiography (ECG) channel for control analysis and artefact removal [12]. While a single channel is normally sufficient to identify heartbeat timings, it cannot fully capture the multidimensional structure of cardiac artefacts and reflects only some projections depending on its location [8, 23]. The cardiac artefact occupies a subspace of the high-dimensional EEG: atrial and ventricular depolarisation, repolarisation, and the ballistocardiographic response each produce topographically and temporally distinct patterns [17, 24, 25]. This complexity is compounded by the rotation of the heart during the cardiac cycle, which causes the artefact topography to vary over time [8]. Identifying this subspace reliably requires a reference signal of sufficient dimensionality, a requirement that a single ECG channel cannot satisfy [23].

The choice of ECG channel placement is an additional complication. Standard limb and chest channels sample the cardiac field at the torso, but the artefact contaminating scalp EEG propagates through the neck and head [23]. Electrodes positioned along this conduction path may better represent the artefact’s spatial structure as it projects toward the scalp and also capture pulse-related movement artefacts.

Canonical correlation analysis (CCA) offers a natural solution to the multi-channel artefact identification problem. By finding linear combinations of EEG channels that maximally correlate with linear combinations of a multi-channel ECG reference, CCA directly identifies the EEG subspace most contaminated by cardiac activity without requiring prior specification of the number of artefact components [26]. Each canonical component corresponds to a direction in EEG sensor space maximally explained by the multi-channel cardiac reference, and components can be ranked by their canonical correlation coefficients, providing a principled ordering from most to least artefact-contaminated. Critically, unlike ICA, which blindly decomposes the EEG and then relies on post hoc comparison with a single ECG channel to identify cardiac components, CCA incorporates multi-channel cardiac information directly into the decomposition, using the full spatial structure of the recorded cardiac field to guide artefact identification.

The practical application of any subspace removal method raises the critical question of how many components to remove [12, 20]. Removing too few components leaves residual artefacts but removing too many risks suppressing genuine neural signals that share spatial or temporal structure with the cardiac reference. This tradeoff is particularly acute for HER research, where the neural signal of interest is time-locked to the R-peak [27]. Here we propose the R-peak amplitude as a principled stopping criterion: the R-peak appears in the EEG signal as a direct consequence of volume conduction at a latency too short for genuine neural processing and therefore it is likely to largely consist of artefact. Monitoring the EEG R-peak amplitude after cleaning as a function of the number of removed components provides a direct and interpretable measure of residual cardiac noise at each component removal step. Thus, one can identify stopping points where no substantial improvement in cleaning efficacy occurs with additional removed components.

In the present study, we systematically compare ICA- and CCA-based cardiac artefact removal using a 15-channel ECG recorded from 14 participants, allowing us to characterize artefact suppression as a function of the number of removed components and the number and placement of ECG channels. We propose the R-peak amplitude as a principled data-driven stopping criterion, and provide recommendations for ECG channel configuration and electrode placement in heartbeat-locked EEG research. Our results have direct implications for the design and reporting of cardiac artefact removal pipelines in interoception research, autonomic neuroscience, and any application requiring disentanglement of cardiac and neural signals.

## 2 Materials and Methods

### 2.1 Experimental Procedure

#### 2.1.1 Ethics

The study was approved by the Ethics Committee of the Medical Faculty at the University of Leipzig (number 280/23-EK) in accordance with the Declaration of Helsinki. Control group participants were recruited by the Max Planck Institute for Human Cognitive and Brain Sciences using the institute’s participant database. Participants received a compensation of 15 euro per hour. Data reported here are part of a larger project also involving patients with heart arrhythmia; only resting-state data from the control group, consisting of 14 participants, are analysed in the present study.

#### 2.1.2 Data Acquisition

Participants underwent a clinical screening visit, during which eligibility was assessed and written informed consent was obtained. Holter ECG monitoring and echocardiography were performed to assess cardiac status and exclude relevant cardiac abnormalities, followed by blood and urine sampling and an MRI safety briefing. Control participants completed a single baseline visit that began with a quality-of-life, neurocognitive, and mood assessment battery. For more details, see Supplementary Information 1. Participants then underwent structural, diffusion, and functional MRI scanning (3 Tesla Siemens Skyra scanner), the latter comprising a resting-state recording and a functional recording during a one-back task; a one-channel II-lead bipolar ECG was recorded concurrently with the fMRI. Later on the same day, a simultaneous EEG-ECG recording was performed, mirroring the fMRI session: it began with two resting-state sessions of 10 minutes each, followed by a task recording using the same one-back paradigm.

#### 2.1.3 Electrophysiological Recording

The EEG recording was conducted in a soundproof, electrically shielded room. EEG signals were recorded with a Brainvision amplifier at a sampling rate of 2500 Hz, bandwidth 0.5–1000 Hz. A 64-channel cap (actiCAP) was used to place electrodes according to a standard 10-20 system. At set-up, electrode impedances were kept at <15 kOhm for all channels. The Fpz electrode served as ground, and FCz served as a reference. The TP9 electrode was used for recording eye movements. 15 ECG channels and one respiration channel were acquired simultaneously with EEG. ECG was recorded using dedicated ECG channels embedded in the BrainProducts montage. ECG included a combination of six standard clinical chest channels (V1–V6) and three standard limb channels (right arm, RA; left arm, LA; left leg, LL) placed in modified torso positions (RA and LA on the right and left upper torso, infraclavicular chest; LL on the lower left torso), augmented by two supraclavicular and four neck electrodes (two anterior, two posterior) to account for blood vessel pulsatility, (see Fig. 1A).

**Figure 1:**
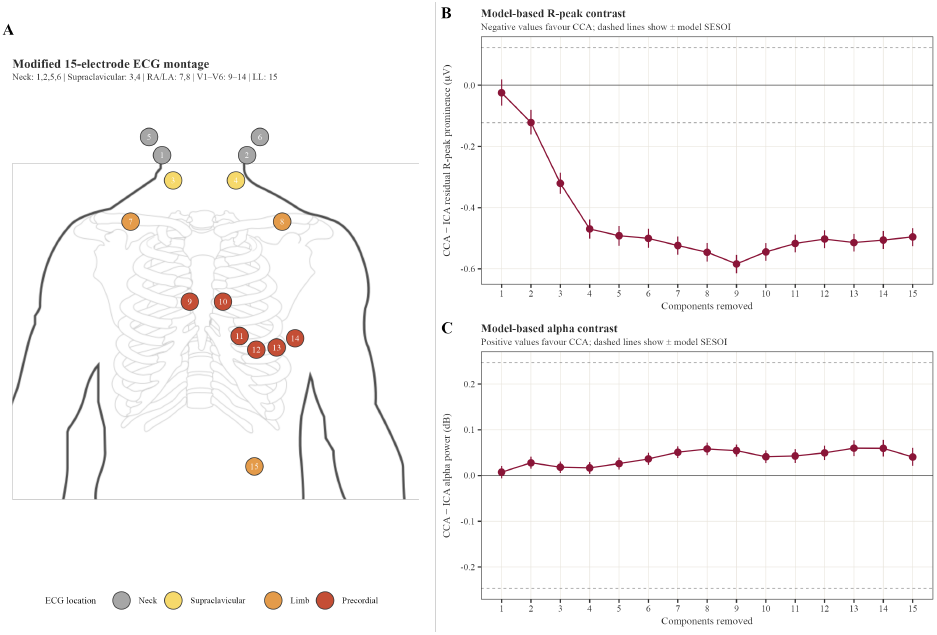
(A) 15-channel ECG setup. Neck electrodes 5 and 6 are on the same level as 1 and 2 but on the back. 3 and 4 are placed above clavicles. 7 and 8 are below clavicles on the chest. 9-14 are standard V1-V6 placements. 15 is on the lower left abdomen. (B) Posterior model estimates for the difference in R-peak prominence and (C) alpha power between ICA and CCA, for different numbers of removed components. Points show posterior medians; vertical bars show 95% posterior intervals. The solid horizontal line marks no difference between methods, and dashed horizontal lines mark the model-based SESOI. ECG - electrocardiography, ICA - independent component analysis, CCA - canonical correlation analysis, SESOI - smallest effect size of interest.

### 2.2 Data Processing

#### 2.2.1 EEG

EEG data were preprocessed in MATLAB using EEGLAB [28] and in-house scripts. After loading the raw data, they were downsampled to 250 Hz and standard electrode locations were assigned using the 10-5 system. Non-EEG channels (ECG, respiration, and EOG) were separated. Line noise at 50 Hz was removed using the CleanLine algorithm [11]. Bad channels were identified and removed in two passes: first using the `clean_rawdata` plugin and second using a variance- and outlier-based rejection procedure applied over 4-second epochs in the band 0.5 to 20 Hz. Data were then re-referenced to the common average and bandpass filtered between 0.5 and 20 Hz using a zero-phase Hamming-windowed sinc FIR filter (pop_eegfiltnew; EEGLAB), with filter order automatically determined based on the specified transition bandwidth.

#### 2.2.2 ICA

ICA was performed using the Infomax algorithm [21], and components were automatically classified with ICLabel [29]. Components reflecting muscle activity or eye movements (probability > 0.8) or channel noise (probability > 0.5) were removed. Finally, previously removed channels were reconstructed via spherical spline interpolation [30], restoring the full channel montage.

#### 2.2.3 ECG, R-peak Detection and HER Epochs

The ECG channel exhibiting the largest peak-to-peak amplitude was selected for R-peak detection. Then, all ECG channels were band-pass filtered between 0.5 and 20 Hz using the same approach as for EEG channels. R-peaks were then detected using the FMRIB EEGLAB plugin [31], which implements a template-based QRS detection algorithm. Detected R-peaks were inserted as discrete event markers and used to epoch both the EEG and ECG signals in a window of −200 to 700 ms relative to each R-peak onset, yielding cardiac-cycle-locked epochs for downstream HER analysis. HER epochs were baseline corrected using -200 to -100 ms pre-R-peak window. No further epoch rejection criteria were applied because both ICA and CCA were applied to continuous EEG or EEG-ECG signals, and HER epochs were only used for visualisation and extraction of EEG R-peak amplitudes (Fig.2).

### 2.3 Canonical Correlation Analysis for Cardiac Artefact Removal

CCA was applied as an alternative to ICA-based removal for separating cardiac field artefact from the EEG signal [32]. CCA decomposes two multivariate signals — here the continuous band-pass filtered (0.5–20 Hz) EEG and ECG — into pairs of canonical variates that are maximally correlated across modalities. Given the EEG data matrix **X** ∈ ℝ ^*T* ×*N*^ with *N* the number of EEG sensors and *T* the number of time samples, and the ECG data matrix **Y** ∈ ℝ ^*T* ×*M*^ with *M* the number of ECG sensors, CCA finds weight vectors **w**_*x*_ and **w**_*y*_ that solve:

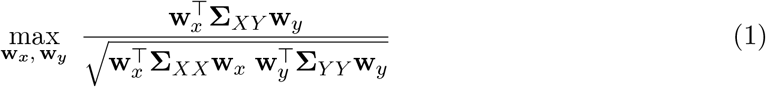

where **∑** _*XX*_ and **∑** _*Y Y*_ are the sample covariance matrices of the EEG and ECG respectively, and **∑** _*XY*_ is their cross-covariance matrix. The full set of spatial filters **W**_*x*_ ∈ ℝ^*N* ×*N*^ and **W**_*y*_ ∈ ℝ^*M* ×*M*^ is obtained by solving Equation 1 successively under the constraint that each pair of canonical variates is uncorrelated with all previous pairs. The corresponding spatial activation patterns were recovered as [33]:

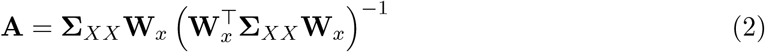

Components were ranked by their canonical correlation coefficient and removed from the EEG iteratively in descending order, up to a maximum of 15 components. At each iteration *k*, let **W**_*x,k*_ and **A**_*k*_ denote the filters and patterns of the first *k* components to be removed. The cardiac artefact subspace was estimated as:

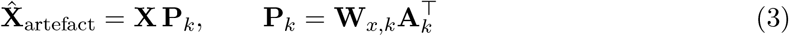

where **P**_*k*_ ∈ ℝ ^*N* ×*N*^ is an oblique projector onto the cardiac subspace spanned by the first *k* canonical filters, constructed such that 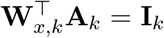 (biorthogonality). The cleaned EEG at iteration *k* was then obtained by subtracting the artefact estimate:

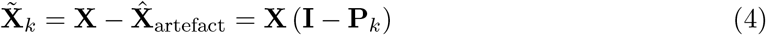

In practice, CCA was computed using the *canoncorr* function of MATLAB, and spatial patterns as well as component removal were estimated using custom code.

### 2.4 Iterative Artefact Removal and Cleaning Efficacy

In this section, we first describe the iterative procedure we use to remove CFA from the EEG. Moreover, since there are currently no widely accepted procedures for evaluating the effectiveness of CFA removal, we propose two complementary approaches, one based on the prominence of the R peak and another on the level of residual alpha oscillations, which are also described here.

#### 2.4.1 Iterative Procedure

ICA and CCA were compared for the iterative removal of CFA from the EEG signal. ICA-based removal was applied by ranking continuous ICA components [32] according to their absolute Pearson correlation with the bandpass-filtered (0.5–20 Hz) continuous ECG signals [34]. Each component was correlated with all ECG channels, and the maximum absolute correlation value was retained. Components were then removed iteratively in descending order of correlation. CCA was applied jointly to the continuous EEG and ECG data, decomposing the EEG into components of maximal correlation with the ECG channels, which were subsequently iteratively removed as described in Section 2.3.

#### 2.4.2 R-peak Prominence

After each removal step (ICA- or CCA-based) the residual CFA was quantified using an R-peak amplitude metric. For each participant, a time window of interest around the R-peak was individually defined by visually inspecting the trial-averaged and baseline corrected EEG across all channels. The mean onset of the R-peak was −57.1 ± 39.9 ms and the mean offset was 96.1 ± 18.3 ms relative to the R peak (mean window duration: 153.2 ± 51.8 ms; full range: -100 to 125 ms across participants, Supplementary Information 2). The maximum absolute amplitude of the trial-averaged signal within this window was then computed per channel and used as an index of residual cardiac artefacts.

#### 2.4.3 Alpha Power

To assess the effect of artefact removal on neural signal integrity, changes in alpha power were evaluated following ICA and CCA component removal. The individual alpha band was first defined per participant from the observation of the power spectral density (PSD) of the continuous preprocessed EEG, averaged over channels and prior to any cardiac component removal. PSD was estimated in the continuous EEG data using the Welch’s method and 4-second segments, yielding a frequency resolution of 0.25 Hz. The segments overlapped 50% and the data was windowed using the default Hamming window. The resulting periodograms were averaged across segments to produce a smoothed PSD estimate for each channel. The baseline alpha power value was obtained by averaging the PSD in the individually selected alpha band in each channel (mean lower bound 8.09 ± 0.95 Hz, mean upper bound, 11.71 ± 1.75 Hz and mean bandwidth 3.63 ± 1.36 Hz, with full-range 6.75–14.75 Hz across participants, Supplementary Information 2). The estimation of alpha power after ICA or CCA component removal was performed by computing the PSD of the continuous residual EEG as described above. The averaged PSD in the individually selected frequency band was compared to the baseline estimate.

#### 2.4.4 Smallest Effect of Interest

To distinguish negligible numerical differences from practically meaningful changes, we defined a smallest effect of interest (SESOI) as 5% of the no-cleaning baseline value. For descriptive summaries, SESOI values were computed separately for each participant and EEG channel. For all-channel model-based contrasts, we used a pooled model SESOI equal to 5% of the mean no-cleaning baseline across participant-by-EEG-channel cells.

### 2.5 Statistical Analyses

All statistical and descriptive analyses were conducted in R. Bayesian multilevel models were fitted using brms [35] with the cmdstanr backend [36]. R-peak prominence and alpha power were analysed in separate models. For both outcomes, the main model included cleaning method, cleaning level, and their interaction as fixed effects, with random intercepts for participant, EEG channel, and participant-by-channel combinations:

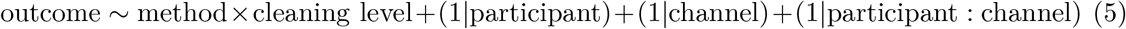

Models were fitted with a Student-(t) likelihood to reduce sensitivity to extreme observations. The cleaning method was coded as ICA or CCA, and the cleaning level was treated as a categorical factor to avoid imposing a linear dose-response assumption across removed components. Models were fitted with four chains, 4000 iterations per chain, 2000 warm-up iterations, adapt_delta = 0.95, and max_treedepth = 12. Convergence was assessed using standard posterior diagnostics, including trace plots and 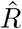.

Posterior marginal means and contrasts were estimated using emmeans [37], and posterior draws were extracted using tidybayes [38]. Results are reported as posterior medians with central 95% posterior intervals (PI), together with posterior probabilities for directional and SESOI-based hypotheses. For method comparisons, contrasts were computed as CCA minus ICA; negative values therefore indicate lower residual R-peak prominence after CCA. For cleaning-level analyses, consecutive contrasts between adjacent cleaning levels were used to assess whether removing an additional component produced a SESOI-meaningful reduction in R-peak prominence and to support model-based stopping-point decisions. For alpha power, the same modelling and contrast logic was used, but effects were interpreted with respect to preservation of neural signal strength.

Additional descriptive summaries of observed values are reported as medians, interquartile or percentile ranges, and percentages of observations falling into SESOI-based categories, including “CCA lower than ICA beyond SESOI”, “practically similar”, and “ICA lower than CCA beyond SESOI”. Equivalent SESOI-based summaries were used for R-peak reduction distributions across cleaning methods, cleaning levels, participants, and channels to complement model-based conclusions due to the high heterogeneity in observations.

### 2.6 Top-Performing ECG Sets and Anatomical Locations

Reduced-set ECG configurations were analysed for CCA only. For each participant and EEG channel, the full 15-channel reference was defined as the lowest residual R-peak prominence achieved across all available CCA cleaning levels. The corresponding no-cleaning value was used to define the participant-by-channel SESOI.

We evaluated all possible ECG combinations containing 1, 2, 3, 4, or 5 channels, corresponding to 15, 105, 455, 1365, and 3003 exact combinations, respectively. For each exact reduced combination, participant, and EEG channel, we selected the optimal cleaning level, defined as the level with the lowest residual R-peak prominence. Reduced combinations were compared with the full 15-channel optimum using:

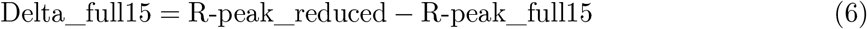

Because a lower R-peak prominence indicates better cleaning, a reduced combination was classified as not meaningfully worse than full 15 when Delta_full15 ≤ SESOI (in our case 5%). Loss was additionally expressed in SESOI units and as a percentage of the no-cleaning baseline.

Exact ECG combinations were ranked separately within each channel count. The primary criterion was participant-balanced EEG-channel coverage: for each participant, we computed the percentage of available EEG channels for which a combination was not meaningfully worse than the full 15-channel set, and then summarised these values across participants. Ranking additionally considered the number of participants with acceptable performance in at least 50% of EEG channels, median and lower-quartile channel coverage, median and upper-tail loss in SESOI units, and the percentage of full 15-channel cleaning improvement recovered. In addition, we inspected aggregated participant-by-EEG-channel metrics to quantify overall performance.

We assessed top-performance robustness using two bootstrap procedures. Participants were resampled with replacement 100 times, and EEG channels were resampled with replacement 100 times; exact ECG combinations were re-ranked in each bootstrap sample. For each channel count, we summarised robustness as the percentage of bootstrap samples in which original top-performing candidates reappeared among high-ranking combinations. Main top-pool summaries were based on the top 20% of combinations.

Anatomical interpretation was performed after exact combinations had been ranked. ECG channels were annotated as neck, supraclavicular, standard limb, or precordial/V locations (Fig.1A). Each exact combination was assigned to one mutually exclusive family: all V/precordial, all neck/supraclavicular, all limb anchors, V + limb, V + neck/supraclavicular, neck/supraclavicular + limb, or V + neck/supraclavicular + limb. We then summarised family prevalence among the top 20% of exact combinations within each ECG-channel count. Thus, exact combination ranking and bootstrap robustness formed the primary result, while anatomical labels were used to summarise recurring spatial principles among robust candidates.

## 3 Results

### 3.1 CCA and ICA Comparison

Fig. 2 (Original row) shows the uncleaned R-peak-locked scalp topography for two participants. For Participant 1, the artefact is asymmetric, with the largest amplitudes (up to 3 *µ*V) over left posterior, occipital and frontal peripheral electrodes. For Participant 2, the topography appears more diffuse, reaching higher absolute amplitudes (up to 6 *µ*V) with only a modest left-lateralised asymmetry. In both participants, the amplitude and spatial extent of the artefact are larger than what would be expected from genuine, spatially focal cortical activity at this latency, and its timing is fixed to the R-peak rather than to any experimental event.

**Figure 2:**
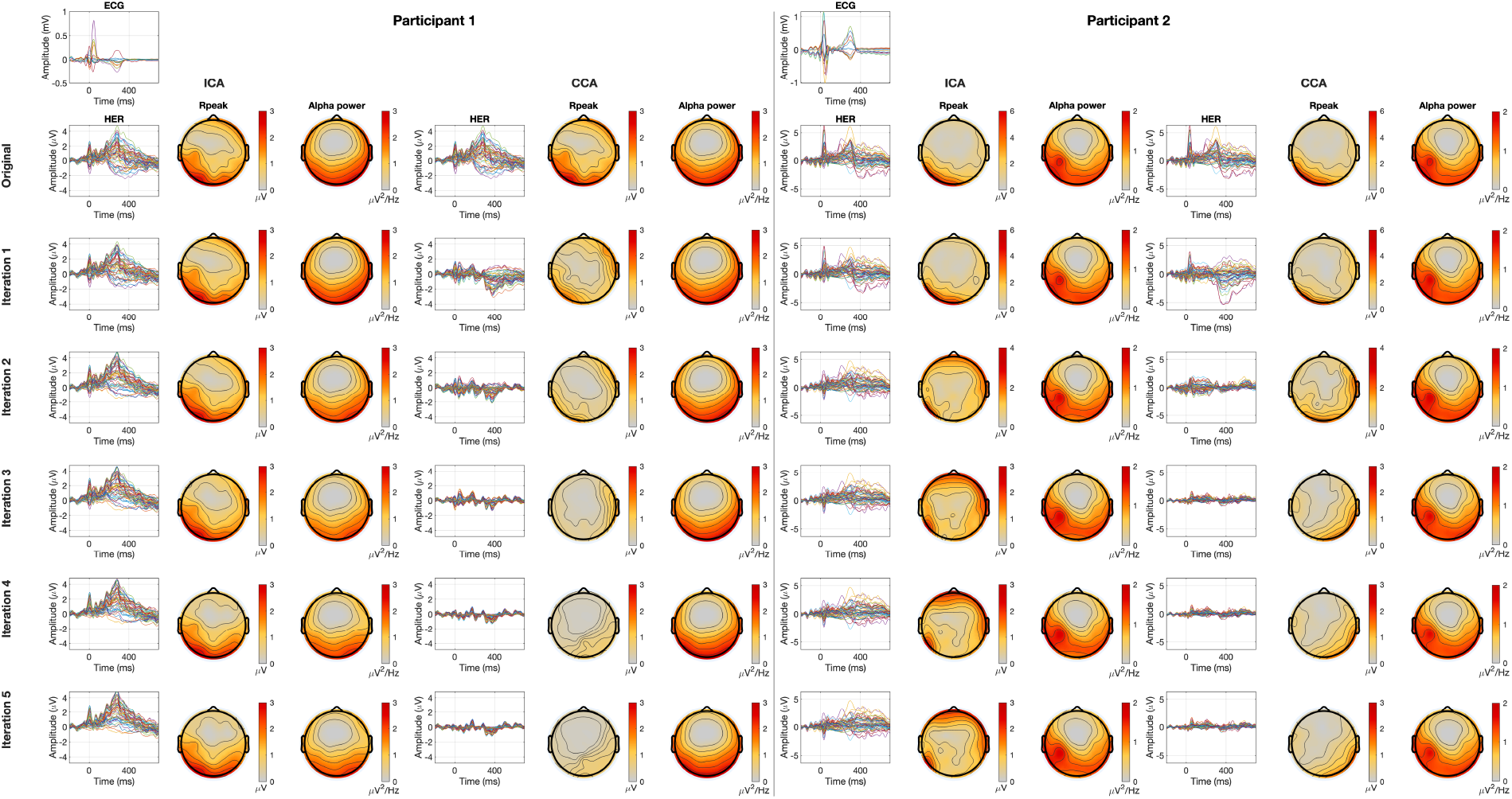
For ICA and CCA separately, averaged ECG and EEG responses locked to R-peaks, as well as R-peak artefact and alpha power topographies across cleaning levels for two participants. ECG - electrocardiography, EEG - electroencephalography, ICA - independent component analysis, CCA - canonical correlation analysis, HER - heartbeat-evoked response.

CCA reduced residual R-peak prominence more effectively than ICA. Across cleaning levels and EEG channels, the posterior contrast CCA minus ICA was negative, indicating lower R-peak artefact after CCA: median difference = -0.416 *µ*V, 95% PI -0.426 to -0.407, P(CCA<ICA) > 0.999; P((CCA-ICA)*<* −SESOI) > 0.999 (Fig. 1B). This model-based result was supported by the observed SESOI-based counts: CCA produced lower R-peak prominence than ICA beyond SESOI in 79.2% of participant-by-channel-by-cleaning-level observations, whereas ICA was lower beyond SESOI in only 10.8%, with 10.0% of observations classified as practically similar.

The advantage of CCA was dependent on cleaning level. After removing only the first component, ICA and CCA showed broadly comparable R-peak reductions: P(CCA<ICA) = 0.872, but P(|CCA-ICA| ≤ SESOI) > 0.999. However, as more components were removed, CCA increasingly outperformed ICA (after three, P(CCA<ICA) > 0.999, P((CCA-ICA)*<* −SESOI) > 0.999), consistent with a more effective use of multi-channel cardiac information. This pattern was visible both in the model-based cleaning-level contrasts and the observed percentages of participant-by-channel observations favouring CCA at later cleaning levels (Supplementary Information 4).

Importantly, the stronger artefact reduction after CCA was not accompanied by stronger loss of alpha power. Across cleaning levels, alpha power after CCA was comparable to, or slightly higher than, alpha power after ICA: median CCA-ICA alpha contrast 0.05 dB, 95% PI 0.04 to 0.05 dB; P(CCA > ICA) > 0.999, P(|CCA-ICA| ≤ SESOI) > 0.999 (Fig. 1C). Observed SESOI-based counts showed that alpha power was lower after CCA beyond SESOI in 30.1% of observations, comparable in 29.8%, and higher after CCA beyond SESOI in 40.1%. Thus, CCA improved cardiac artefact suppression while preserving alpha power at least as well as ICA (Fig. 2).

### 3.2 Stopping Point Analysis

Cleaning-level analyses showed different stopping-point profiles for ICA and CCA. For both methods, the removal of the first component produced the clearest reduction in R-peak prominence (ICA: 56.3% observations with SESOI-meaningful decrease; CCA: 61.3%; model probability of SESOI-meaningful decrease: 0.999 and 1.000). Beyond this first component, ICA showed less consistent improvement: some participant-by-channel observations continued to improve at subsequent cleaning levels, but these reductions were sporadic and did not form a stable gradual pattern across participants and channels. Consequently, model-based consecutive contrasts did not support a clear ICA stopping point beyond the first removed component: after the second component, the probability of a meaningful decrease was only 0.001.

By contrast, CCA showed a more gradual and reproducible reduction in R-peak prominence across consecutive cleaning levels. SESOI-meaningful decreases were still evident after four removed components. For added components 2–4, observation-level SESOI-meaningful decreases occurred in 56.5%, 68.3%, and 55.1% of participant-by-channel observations, respectively; the corresponding model probabilities of SESOI-meaningful decrease were 0.943, 1.000, and 1.000. However, the probability and magnitude of additional improvement decreased thereafter. Across model-based contrasts and observed stopping-level distributions, CCA showed a practical plateau at around four components: the median number of SESOI-meaningful improvements was 4, and 77.0% of participant-by-channel observations showed 3-5 SESOI-meaningful component removal steps.

Observation-level stopping summaries were essential because stopping behaviour was heterogeneous. Under a conservative rule, defined as stopping at the first non-useful component, ICA stopping points were concentrated early, with only 4.9% of participant-EEG-channel observations falling in the 3–5 component range. Under a more liberal rule, defined by the last SESOI-meaningful improvement, ICA frequently extended to late cleaning levels, including the maximum level in 7.6% of participant-by-channel observations; 68.5% of observations extended to ten or more removed components. CCA was less extreme under the liberal rule: the median retained step after the last useful improvement was 9 and the mode was 4 (Supplementary Information 5). Thus, ICA produced either early or very late stopping estimates depending on the rule, whereas CCA showed a more stable plateau (Fig. 3A).

**Figure 3:**
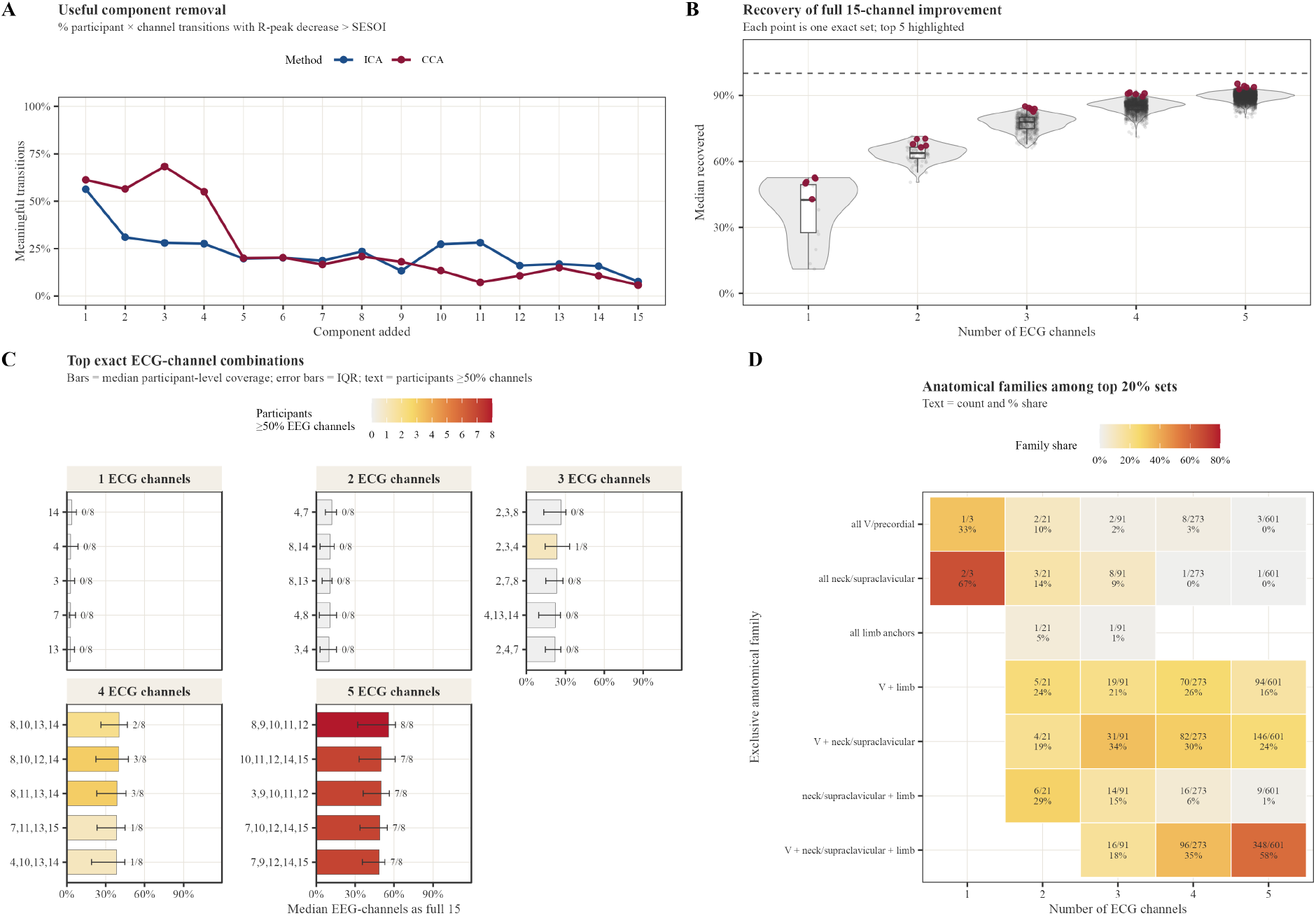
Percentage of participant-EEG-channel transitions in which removing the next component produced a meaningful R-peak decrease larger than the SESOI, separately for ICA and CCA. (B) Recovery of the full 15-channel CCA improvement by reduced ECG-channel configurations. Each grey point is one exact ECG-channel combination. Violin plots show the distribution of median recovered full-15 improvement across exact combinations for each ECG-channel count; white boxplots show the median and IQR. Bordeaux points highlight the five top-ranked exact CCA combinations within each ECG-channel count. The dashed horizontal line marks 100% recovery of the full 15-channel CCA improvement. (C) Top exact reduced ECG-channel combinations within each channel count. Bars show the median percentage of EEG channels for which the reduced configuration was not meaningfully worse than full 15-channel CCA. Error bars show the interquartile range across participants. Text labels show the number of participants for whom the configuration reached at least 50% EEG-channel coverage as good as full 15. Fill intensity encodes the same participant-support count. (D) Anatomical composition of high-ranking exact configurations. Tiles show the share of exact top-20% configurations belonging to each mutually exclusive anatomical family, separately for each ECG-channel count. Text gives the count of top configurations in that family and the corresponding family share. Fill intensity encodes family share. ECG - electrocardiography, EEG - electroencephalography, ICA - independent component analysis, CCA - canonical correlation analysis, SESOI - smallest effect size of interest, IQR - interquartile range.

### 3.3 Preferred ECG Combinations

Reduced-channel analyses evaluated whether subsets of the 15 ECG channels could reproduce the best full-montage CCA result. The best one-channel configuration was channel 14 (V6), closely followed by 3 and 4 (left and right supraclavicular). However, it was not consistently comparable to the full 15-channel reference: it was not meaningfully worse than full 15 only in 4.93% of participant-by-EEG-channel observations, with median loss 8.49 SESOI units and 49.98% of the full 15-channel improvement recovered. Thus, a single ECG channel could reduce cardiac artefact substantially in some cases, but did not provide a robust substitute for multi-channel ECG (Fig. 3B).

Additional ECG channels improved performance. The best two- and three-channel configurations were 4,7 (followed by 8, 14 and 8,13) and 2,3,8 (followed by 2,3,4 and 2,7,8), recovering 66.42% and 85.05% of the full 15-channel improvement, respectively, and were not meaningfully worse than full 15 in 12.39% and 23.24% of participant-by-EEG-channel observations. Median loss decreased from 8.49 SESOI units for the best one-channel configuration to 5.16 and 2.50 SESOI units for the best two- and three-channel configurations.

Closer-to-full performance required four or five ECG channels. The best four-channel configuration, 8,10,13,14 (LA,V2,V5,V6), was not meaningfully worse than full 15 in 37.04% of participant-by-EEG-channel observations and reached >=50% EEG-channel as-good-as-full-15 coverage in 2/14 participants, with median loss 1.53 SESOI units and 90.49% of the full 15-channel improvement recovered. The best five-channel configuration, 8,9,10,11,12 (LA,V1,V2,V3,V4), performed better: 49.58% of observations were not meaningfully worse than full 15, it reached >=50% EEG-channel coverage in 8/14 participants, median loss was 0.85 SESOI units, and 95.28% of the full 15-channel improvement was recovered (Fig. 3C).

The anatomical composition of top configurations changed with ECG-channel count. For sparse configurations, top-20% pools often included neck/supraclavicular and/or limb-anchor channels. For four and five channels, high-ranking configurations were consistently V/precordial-containing, but not exclusively precordial: the by far dominant five-channel top-20% family of sets combined V/precordial, neck/supraclavicular, and limb-anchor channels (Supplementary Information 6). Note that, with any ECG setup, ICA-based cleaning is unable to provide performance comparable to the full-15 CCA (Supplementary Information 7).

The mutually exclusive anatomical-family analysis quantified this pattern (Fig. 3D). Among top-20% configurations, the dominant family was all neck/supraclavicular for one-channel sets (2/3; 66.7%), neck/supraclavicular + limb for two-channel sets (6/21; 28.6%), V + neck/supraclavicular for three-channel sets (31/91; 34.1%), and V + supraclavicular + limb for four- and five-channel sets (96/273; 35.2%, and 348/601; 57.9%, respectively). In recurring subsets of top 20% 5-channel combinations, there was no strong preference observed for RA, LA, or LL, although LA occurred more frequently (40.8% of all top 5-channel sets) compared to LL (34.3%) or RA (32.9%). Thus, sparse ECG cleaning benefits from upper thoracic/neck-adjacent and limb-anchor information, whereas reduced montages approaching full 15-channel CCA increasingly rely on V/precordial channels embedded in broader mixed anatomical configurations.

## 4 Discussion

Heartbeat-locked EEG analyses face a central methodological problem: the signal of interest is temporally aligned with the cardiac event that also generates the artefact. This makes it difficult to distinguish genuine heartbeat-evoked neural activity from cardiac field artefacts, pulse-related movement, and other physiological signals that covary with the heartbeat. The development of transparent and effective cleaning approaches is therefore essential for heart-brain research [12]. In the present study, we address this problem in three ways. First, we introduce Canonical Correlation Analysis (CCA) as a practical method for integrating multi-channel ECG information into cardiac artefact removal. Second, we propose residual R-peak prominence and preserved alpha power as data-driven criteria for the evaluation of cleaning efficacy and the determination of stopping points. Third, we argue that the heartbeat-evoked responses field should more carefully account for the multivariate nature of cardiac artefacts.

### 4.1 Multi-channel CCA as a Principled Cleaning Method

CCA is a well-known technique and has been widely used in neuroscience and signal processing [26]. Here, we show it can be used as a principled way to integrate multi-channel ECG information into EEG artefact cleaning. Our results show that CCA reduced R-peak prominence more effectively than ICA while preserving alpha power to a comparable or greater extent. ICA showed a strong effect after removing the first component, but less consistent improvement than CCA when removing additional components. This likely reflects that ICA is a blind source separation method and does not guarantee that individual components isolate cardiac from neuronal activity. Rather, the CFA might be spread across many components, leading to unstable artefact suppression. In contrast, CCA explicitly identifies EEG components that covary with multi-channel cardiac activity. This leads to a more gradual reduction in residual R-peak prominence and a clearer plateau after four removed components.

The reduced-channel analyses further suggest that the dimensionality and placement of the ECG reference matter. A single ECG channel was not sufficient to reproduce the full 15-channel CCA solution reliably, even when optimally placed. When only one to three ECG channels were available, neck and supraclavicular locations tended to perform best, in combination with standard limb electrodes. This may be because these locations lie closer to the volume-conducted path from heart to scalp and may also capture vascular or pulse-related activity relevant for scalp artefacts. However, with four or five ECG channels, the best-performing configurations were dominated by precordial/V electrodes, often combined with standard limb or supraclavicular locations. This suggests that once enough electrodes are available, sampling the cardiac field more directly across the chest becomes more valuable than relying only on upper thoracic or neck-adjacent locations. Furthermore, among 5-channel sets, combinations of precordial, limb, and supraclavicular locations performed better than exclusively precordial sets. This indicates that not only information about the heart movement, but also the field spread and possibly vessel pulsations should be integrated to improve cleaning performance.

Practically, these findings support recording as many ECG channels as feasible when heartbeat-locked analyses are planned. If full multi-channel ECG is not possible, a reduced montage with four to five electrodes appears to be the most promising compromise, prioritising several precordial/V locations and at least one limb or supraclavicular electrode, but ideally both. If even fewer sensors are available, supraclavicular electrodes may be preferable to arbitrary single-channel placements, especially when the goal is artefact cleaning rather than only R-peak detection. These recommendations do not imply that fewer ECG channels are always adequate for physiological interpretation. Performance varied across participants and EEG channels, and bootstrap analyses confirmed that this variability was largely driven by participant-level differences in the best-performing channels. Efficient artefact cleaning may be achievable with a reduced montage, but condition- or behaviour-related differences in cardiac activity may appear differently across ECG leads [23]. Therefore, richer ECG recordings remain preferable. Even after cleaning, heartbeat-locked analyses should control for heart rate and other relevant cardiac or respiratory parameters [12].

### 4.2 R-peak Prominence and Alpha Power as Cleaning Criteria

A method for artefact removal is only as useful as the criteria available to evaluate it. Currently no widely accepted procedures exist for assessing the effectiveness of cardiac artefact cleaning or for determining how many components to remove. We propose two complementary criteria that together address this gap: residual R-peak prominence as a measure of remaining artefact, and alpha power as a measure of preserved neural signal.

R-peak prominence provides a useful and conservative index of residual cardiac artefact. The R-peak appears in EEG at a latency too short to plausibly reflect cortical processing and therefore offers a direct measure of residual electrical cardiac contamination. However, this criterion is not exhaustive. It targets the direct cardiac field artefact most clearly, but delayed pulse-related artefacts and neural responses may overlap in later time windows. Thus, strong R-peak reduction increases confidence that direct cardiac artefact has been suppressed, but it does not exhaustively prove that all heartbeat-locked non-neural activity has been removed. Alpha power complements R-peak prominence by providing a positive control: a decrease in alpha power possibly indicates that cleaning has removed genuine neural activity rather than artefact alone. However, stable alpha power does not guarantee that no neural activity was removed, since a decrease in alpha power is a sufficient but not necessary marker of neural signal integrity.

In practice, these criteria define a stopping rule, where one monitors whether R-peak prominence decreases beyond a meaningful threshold and whether alpha power remains stable. When further removal no longer meaningfully improves artefact suppression, the cleaning should stop. The present data suggests that approximately three to five removed CCA components may be a reasonable practical range. However, we do not propose this as a fixed cutoff. Stopping points may vary across recordings, participants, EEG systems, artefact magnitude, and MEG versus EEG.

The residual artefact also has implications beyond heartbeat-locked research. If a residual, broadband cardiac artefact can still be detected via R-peak prominence after standard ICA cleaning, as we show here, then this same residual is likely also present in the aperiodic part of the EEG spectrum. In fact, aperiodic (1/f) EEG parameters, increasingly used as markers of cortical excitation-inhibition balance, appear to be systematically biased by cardiac artefact even after conventional artefact rejection [15, 16]. Residual CFA could easily be mistaken for genuine individual or group differences in aperiodic activity. More broadly, we suggest that studies estimating aperiodic offset or exponent from EEG, particularly when comparing across age groups, clinical populations, or other groups that may also differ in cardiac physiology, would benefit from a simple diagnostic such as the R-peak-prominence criterion before drawing conclusions about cortical changes.

### 4.3 Implications for the Heartbeat-Evoked Response

Together, the proposed CCA cleaning method and evaluation criteria have broader consequences for how heartbeat-evoked responses are studied and interpreted. Cardiac field artefacts are not a single, spatially fixed contamination: complex cardiac activity leads to artefacts that vary in space and time. This multivariate structure means that single-lead artefact removal, whether by regression or by ICA, is unlikely to capture the full artefact subspace. When residual artefact remains in the data, it can mimic or mask genuine neural effects, making it difficult to determine whether differences between experimental conditions reflect brain activity or cardiac contamination.

Our results show that CCA multi-channel cleaning is more conservative than traditional one-channel ICA. This means that CCA may remove components that contain some genuine heartbeat-related neural activity, especially when that activity is strongly coupled to cardiac timing. However, for studies aiming to make claims about cortical heartbeat responses, a conservative cleaning strategy may be preferable to leaving strong residual cardiac field artefacts in the data. Our observations also raise the possibility that the artefact-free HER is smaller and has a higher frequency than often assumed. After cleaning, the remaining heartbeat-locked activity may be faster, weaker, or more spatially specific than the broad low-frequency waveforms currently interpreted as HERs [27]. This should be tested directly in larger datasets.

Both CCA and ICA can remove too little or too much artefact or neural signal. We recommend that researchers evaluate the stability of observed effects across cleaning strengths before drawing conclusions. This is especially relevant for slow heartbeat-evoked components, which overlap spectrally and temporally with cardiac and vascular artefacts. Low-frequency HER effects may be more vulnerable to over-cleaning or to residual artefact confounding. Higher-frequency neural features may be more robust to aggressive cleaning, but they should not be assumed to be immune. We therefore recommend reporting whether key condition or group effects remain stable across plausible cleaning levels [12]. Effects that disappear or reverse with small changes in cleaning strength should be flagged as cleaning-sensitive rather than interpreted as unambiguous neural signals.

### 4.4 Limitations

Several limitations should be noted. The present sample is modest, and the recommended channel configurations require validation in independent EEG and MEG datasets, different recording systems, and different participant populations. The optimal number of CCA components and ECG sensors should not be interpreted as universal cutoff values. They are empirical estimates from the present recording configuration. Future work will benefit from larger open datasets with multi-lead ECG, allowing researchers to test whether similar anatomical principles generalize across laboratories. We also focused primarily on R-peak artefact reduction and alpha power preservation; future studies should assess broader spectral and spatial consequences of CCA cleaning.

## 5 Conclusion

In summary, multi-channel CCA provides a convenient and effective way to use ECG information for cardiac artefact removal in heartbeat-locked EEG analyses. Compared with ICA, it offers a more direct use of cardiac reference signals, stronger R-peak reduction, comparable preservation of alpha power, and a more interpretable cleaning trajectory. Our results suggest that four to five ECG channels, especially precordial electrodes combined with a limb reference, can approach full 15-channel performance, whereas fewer sensors provide only partial and less robust cleaning. More broadly, we recommend that heartbeat-locked studies record richer ECG whenever possible, evaluate multiple cleaning strengths, and treat cleaning-sensitive effects with caution.

## Data Availability Statement

The aggregated, anonymised data supporting the findings of this study together with reproducible markdown files and model fit summaries are available at https://osf.io/ehsrj.

## Code Availability Statement

The analysis codes are available at https://github.com/mariaazanova/CardiCCAClean.

## Acknowledgements

This project was supported by the Max Planck Society. Grants PID2020-118829RB-I00, PID2024-161502OB-I00 and IKERBASQUE supported CV.

## Author Contributions

C.V., M.A.: conceptualisation, methodology, software, validation, formal analysis, investigation, visualisation, data curation, writing – original draft, writing – review and editing, project administration. K.G.: conceptualisation, data curation, writing – original draft, writing – review and editing. E.G.: data curation, writing – review and editing. T.P.S.: writing – original draft, visualisation, writing – review and editing. U.L., T.U.: resources, conceptualisation. A.V.: resources, conceptualisation, funding acquisition. V.V.N.: conceptualisation, methodology, writing – review and editing, supervision.

