## Supplementary Information for "Canonical Correlation Analysis and Multi-Channel Cardiography Improve Artefact Cleaning in Heartbeat-Locked Analyses"

#### 1. Neurocognitive assessment battery

Control participants completed a single baseline visit comprising a quality-of-life and a comprehensive neurocognitive and mood assessment battery. Quality of life was assessed using the SF-36 questionnaire [1,2]. Control participants completed a single baseline visit, which began with a neurocognitive assessment comprising the CERAD battery (Consortium to Establish a Registry for Alzheimer's Disease) [3,4], Trail Making Test A & B [5], MoCA (Montreal Cognitive Assessment) [6], CVLT (California Verbal Learning Test) [7,8], RWT (Regensburg Word Fluency Test) [9], WMS (Wechsler Memory Scale) [10], AES (Apathy Evaluation Scale) [11,12], HADS (Hospital Anxiety and Depression Scale) [13,14], and GHB-56 (General Health Behavior Scale) [15].

### 2. Individual R-peak prominence and alpha power regions

*Table S1.* R-peak time window per participant.

| Participant | Onset (ms) | Offset (ms) | Duration (ms) |
|---|---|---|---|
| P1 | -100 | 100 | 200 |
| P2 | -100 | 125 | 225 |
| P3 | -100 | 100 | 200 |

| | | | |
|---|---|---|---|
| P4 | -100 | 125 | 225 |
| P5 | -100 | 85 | 185 |
| P6 | -100 | 100 | 200 |
| P7 | -10 | 100 | 110 |
| P8 | -30 | 70 | 100 |
| P9 | -10 | 120 | 130 |
| P10 | -20 | 90 | 110 |
| P11 | -40 | 80 | 120 |
| P12 | -50 | 100 | 150 |
| P13 | -20 | 70 | 90 |
| P14 | -20 | 80 | 100 |

*Table S2.* Individually selected alpha band width.

| Participant | Lower bound (Hz) | Upper bound (Hz) |
|---|---|---|
| P1 | 6.75 | 8.75 |
| P2 | 7.5 | 12.5 |
| P3 | 8.0 | 8.75 |
| P4 | 8.5 | 11.0 |
| P5 | 6.75 | 10.0 |
| P6 | 10.0 | 12.5 |
| P7 | 7.0 | 10.0 |
| P8 | 8.0 | 12.0 |
| P9 | 8.5 | 13.0 |
| P10 | 8.5 | 12.0 |
| P11 | 8.25 | 13.75 |
| P12 | 9.0 | 13.0 |
| P13 | 7.75 | 12.0 |
| P14 | 8.75 | 14.75 |

### 3. CCA performance details

*Figure S1.* For two participants, CCA ECG weights, patterns and times series, as well as R of each component.

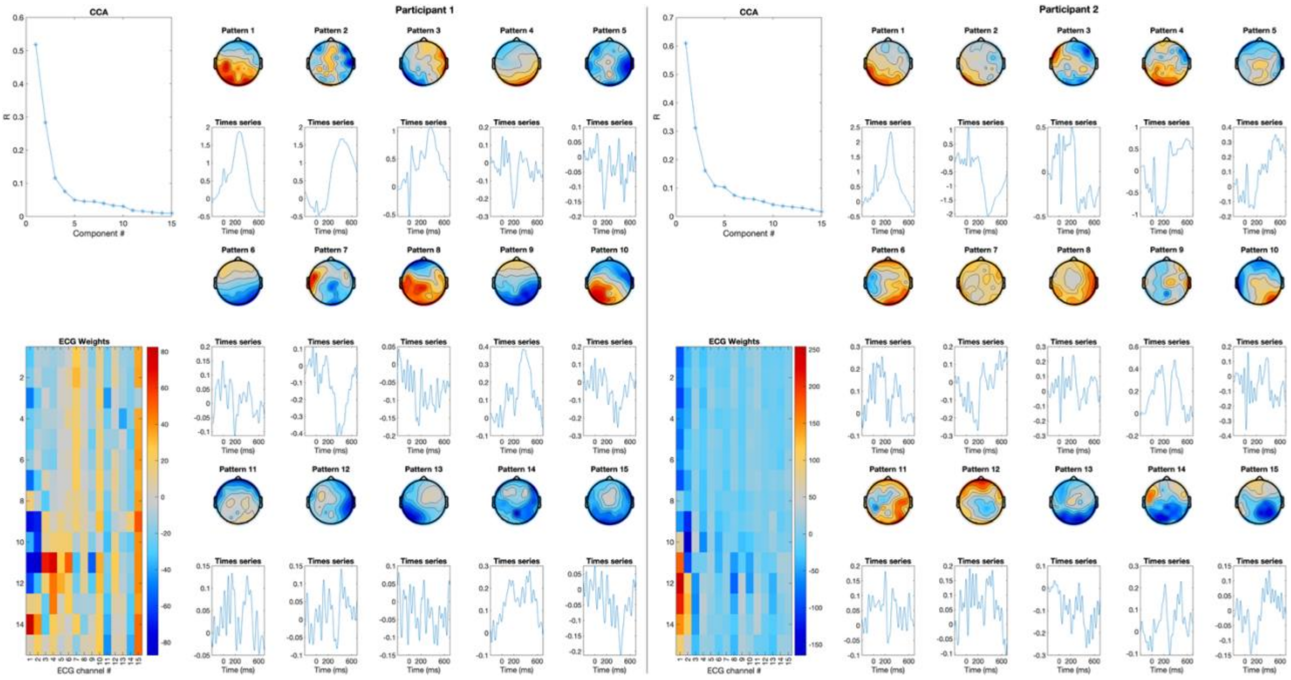


### 4. ICA vs CCA comparisons

*Table S3.* Overall ICA vs CCA comparison for residual R-peak prominence and alpha power.

| **Measure** | **CCA better >SESOI** | **Similar within SESOI** | **ICA better >SESOI** | **Observed median Δ** | **Model median Δ [95%]** | **Model probability** |
|---|---|---|---|---|---|---|
| R-peak, lower better | 79.2% | 10.0% | 10.8% | -0.439 | -0.416 [-0.426, -0.407] | P(CCA lower)=1.000; P(>SESOI)=1.000 |
| Alpha power, higher better | 40.1% | 29.8% | 30.1% | +0.012 | +0.037 [0.033, 0.041] | P(CCA higher)=1.000; P(within SESOI)=1.000 |

*Table S4.* Observed ICA vs CCA comparison by removed component.

| **Comp. removed** | **R-peak CCA/same/ICA %** | **R-peak median Δ** | **Alpha CCA/same/ICA %** | **Alpha median Δ** |
|---|---|---|---|---|
| 1 | 43.4/12.4/44.2 | 0.00 | 15.8/74.7/9.6 | 0.00 |
| 2 | 47.0/11.7/41.3 | -0.05 | 27.8/64.1/8.2 | 0.02 |
| 3 | 68.6/14.8/16.6 | -0.29 | 31.0/56.3/12.7 | 0.01 |
| 4 | 79.7/11.6/8.7 | -0.43 | 30.9/38.3/30.9 | 0.00 |
| 5 | 81.7/8.4/9.9 | -0.49 | 37.6/29.6/32.8 | 0.00 |

| Comp. removed | R-peak CCA/same/ICA % | R-peak median Δ | Alpha CCA/same/ICA % | Alpha median Δ |
|---|---|---|---|---|
| 6 | 81.0/11.4/7.6 | -0.48 | 39.4/24.4/36.2 | 0.00 |
| 7 | 84.4/9.6/6.1 | -0.53 | 44.4/21.6/34.1 | 0.02 |
| 8 | 86.9/8.6/4.5 | -0.60 | 47.3/20.9/31.8 | 0.04 |
| 9 | 88.6/7.8/3.7 | -0.65 | 45.2/22.8/32.0 | 0.03 |
| 10 | 89.2/8.2/2.7 | -0.58 | 44.2/21.7/34.1 | 0.02 |
| 11 | 87.8/8.9/3.4 | -0.53 | 48.0/16.1/35.9 | 0.03 |
| 12 | 87.6/9.0/3.4 | -0.52 | 47.0/16.3/36.6 | 0.03 |
| 13 | 87.8/9.0/3.2 | -0.51 | 48.3/13.8/37.9 | 0.03 |
| 14 | 86.8/10.3/3.0 | -0.50 | 48.7/13.1/38.2 | 0.03 |
| 15 | 87.0/9.2/3.8 | -0.48 | 46.2/13.2/40.6 | 0.01 |

*Table S5.* Model-based ICA vs CCA comparison by removed component.

| Comp. removed | R-peak Δ [95%] | P R-peak >SESOI | Alpha Δ [95%] | P alpha CCA higher | P alpha within SESOI |
|---|---|---|---|---|---|
| 1 | -0.025 [-0.067, 0.019] | 0.000 | 0.007 [-0.006, 0.021] | 0.852 | 1.000 |
| 2 | -0.122 [-0.161, -0.081] | 0.487 | 0.028 [0.015, 0.041] | 1.000 | 1.000 |
| 3 | -0.321 [-0.355, -0.286] | 1.000 | 0.018 [0.005, 0.031] | 0.998 | 1.000 |
| 4 | -0.470 [-0.501, -0.439] | 1.000 | 0.017 [0.004, 0.029] | 0.994 | 1.000 |
| 5 | -0.492 [-0.525, -0.460] | 1.000 | 0.026 [0.013, 0.039] | 1.000 | 1.000 |
| 6 | -0.500 [-0.531, -0.469] | 1.000 | 0.036 [0.024, 0.050] | 1.000 | 1.000 |
| 7 | -0.524 [-0.554, -0.494] | 1.000 | 0.051 [0.038, 0.064] | 1.000 | 1.000 |
| 8 | -0.546 [-0.576, -0.516] | 1.000 | 0.058 [0.045, 0.072] | 1.000 | 1.000 |
| 9 | -0.584 [-0.615, -0.554] | 1.000 | 0.054 [0.041, 0.068] | 1.000 | 1.000 |

| Comp. removed | R-peak Δ [95%] | P R-peak >SESOI | Alpha Δ [95%] | P alpha CCA higher | P alpha within SESOI |
|---|---|---|---|---|---|
| 10 | -0.545 [-0.574, -0.516] | 1.000 | 0.041 [0.027, 0.055] | 1.000 | 1.000 |
| 11 | -0.517 [-0.546, -0.488] | 1.000 | 0.043 [0.027, 0.058] | 1.000 | 1.000 |
| 12 | -0.503 [-0.533, -0.474] | 1.000 | 0.050 [0.034, 0.065] | 1.000 | 1.000 |
| 13 | -0.514 [-0.543, -0.486] | 1.000 | 0.060 [0.042, 0.077] | 1.000 | 1.000 |
| 14 | -0.506 [-0.535, -0.476] | 1.000 | 0.060 [0.042, 0.078] | 1.000 | 1.000 |
| 15 | -0.496 [-0.526, -0.467] | 1.000 | 0.041 [0.021, 0.060] | 1.000 | 1.000 |

*Figure S2.* R-peak prominence after ICA and CCA cleaning.

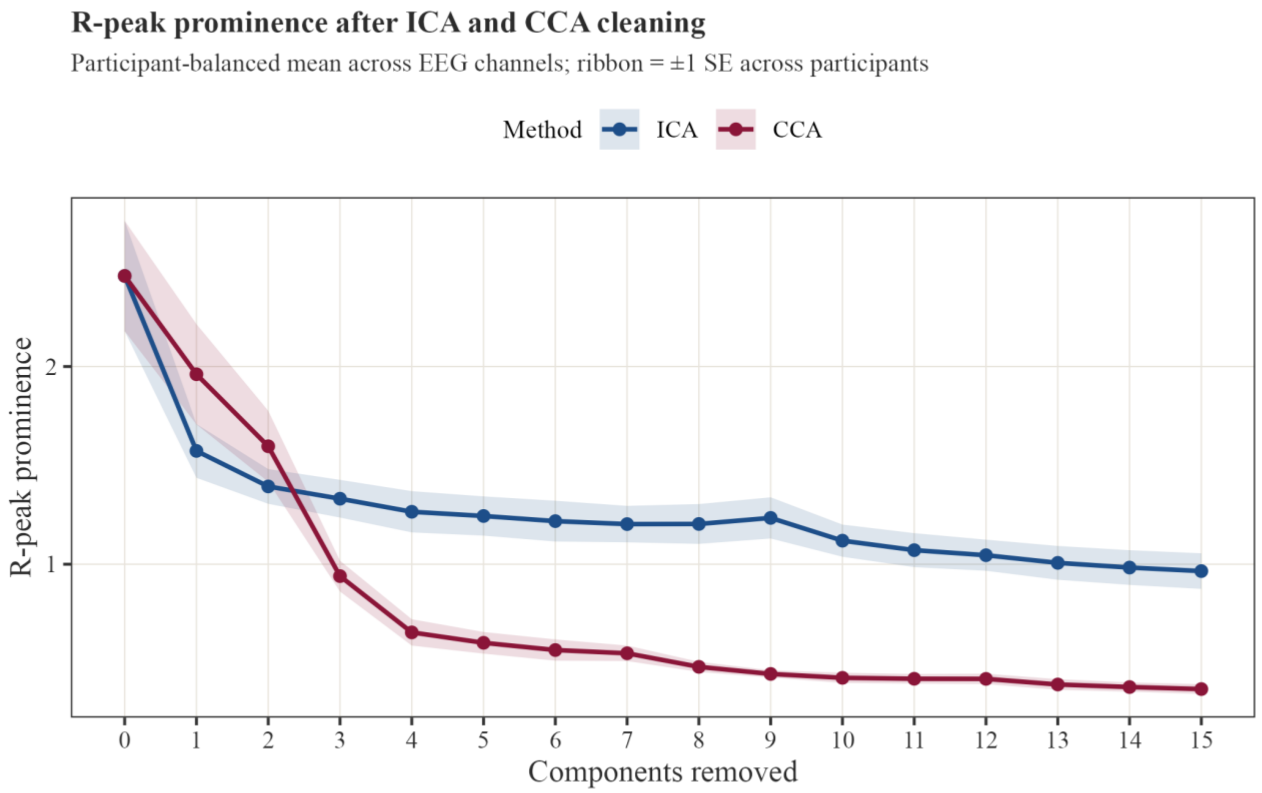


*Figure S3.* Alpha power after ICA and CCA cleaning.

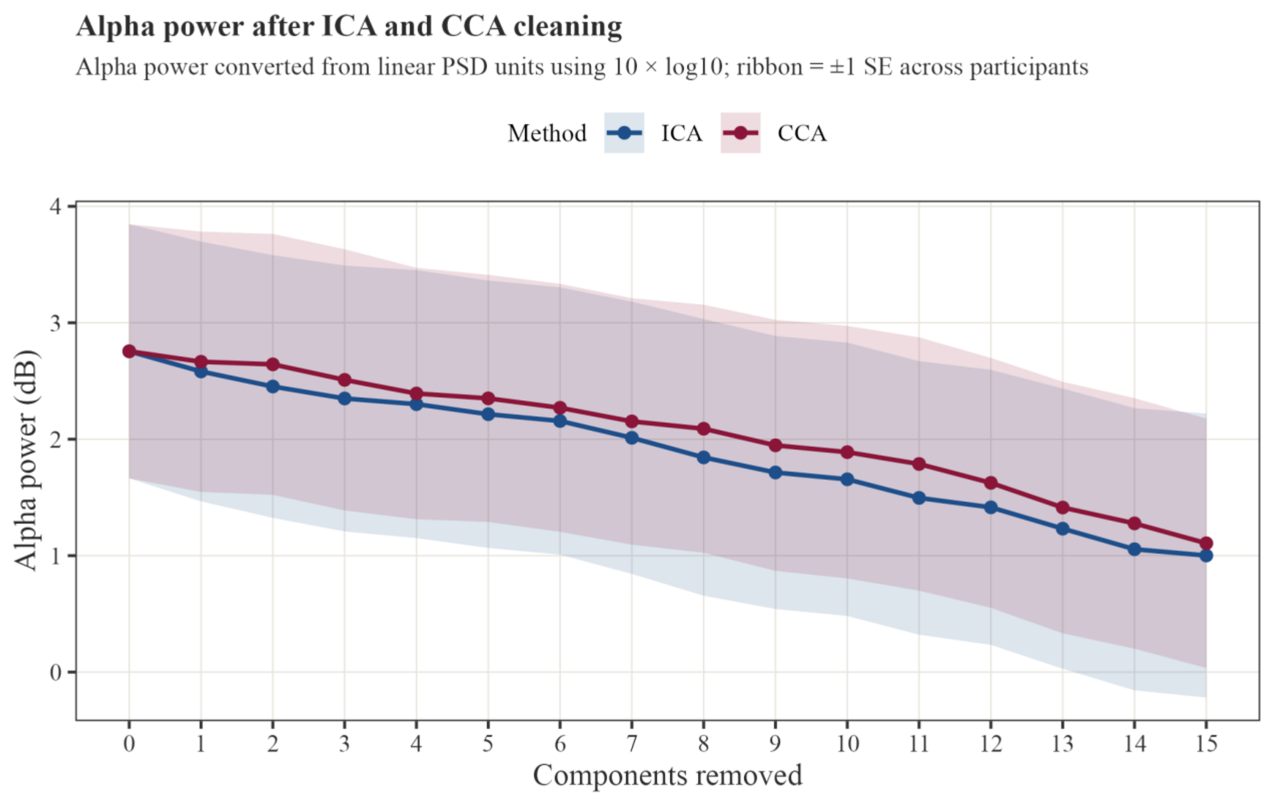


*Figure S4.* Observed ICA vs CCA classifications for R-peak prominence by removed component.

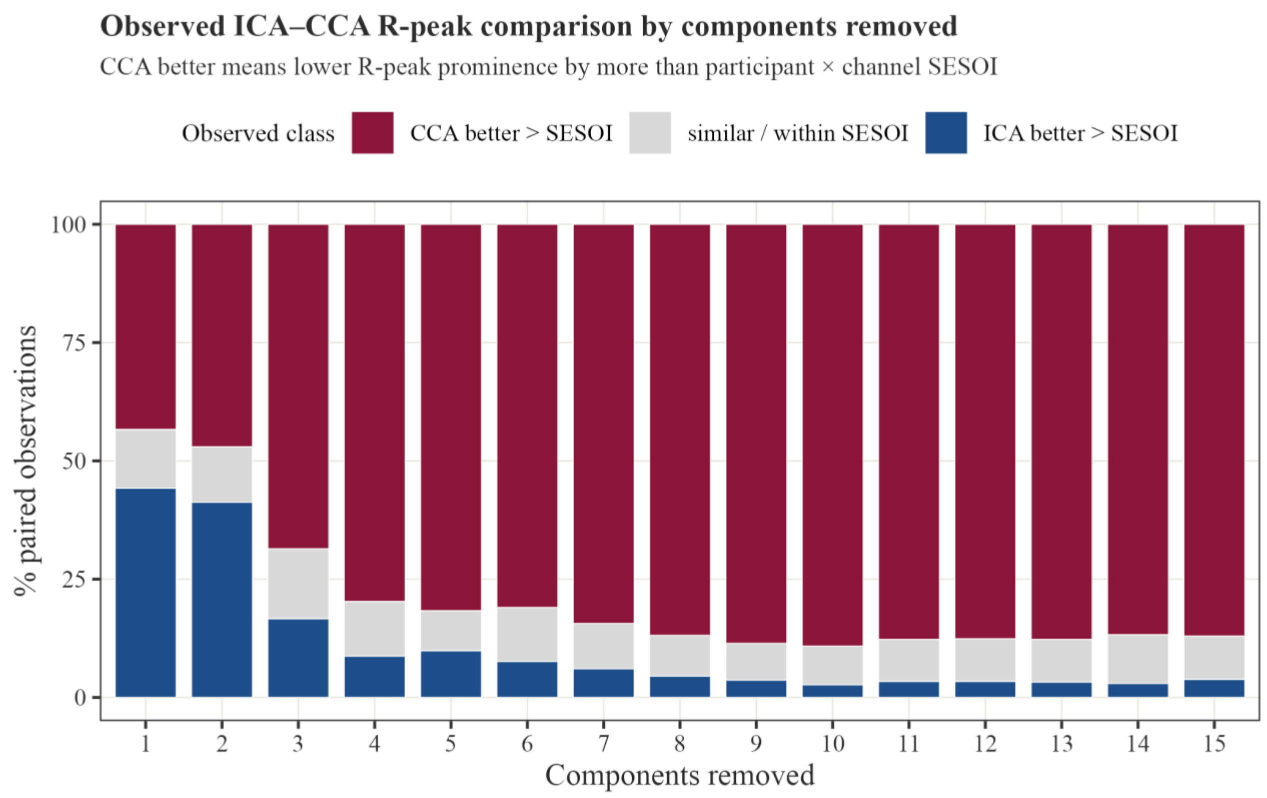


*Figure S5.* Observed ICA vs CCA classifications for alpha power by removed component.

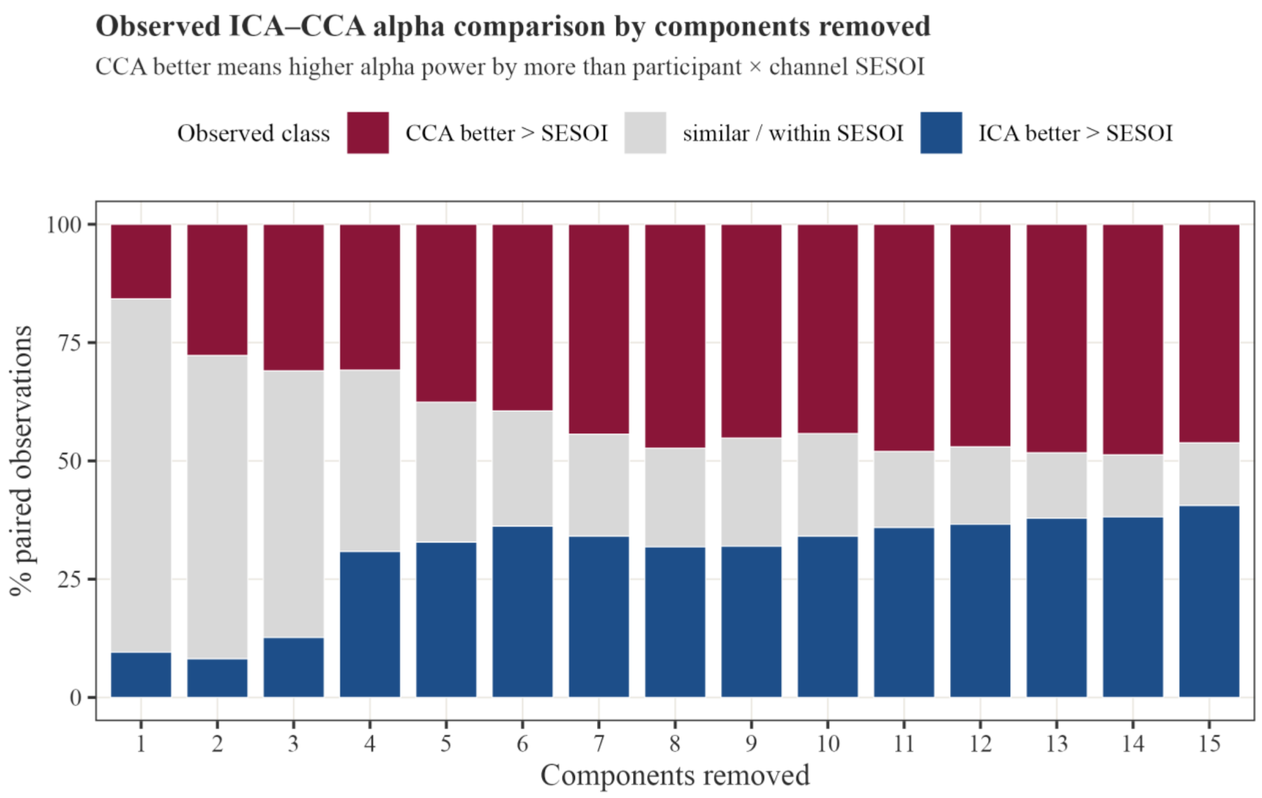


*Figure S6.* Posterior CCA-ICA contrast for residual R-peak prominence.

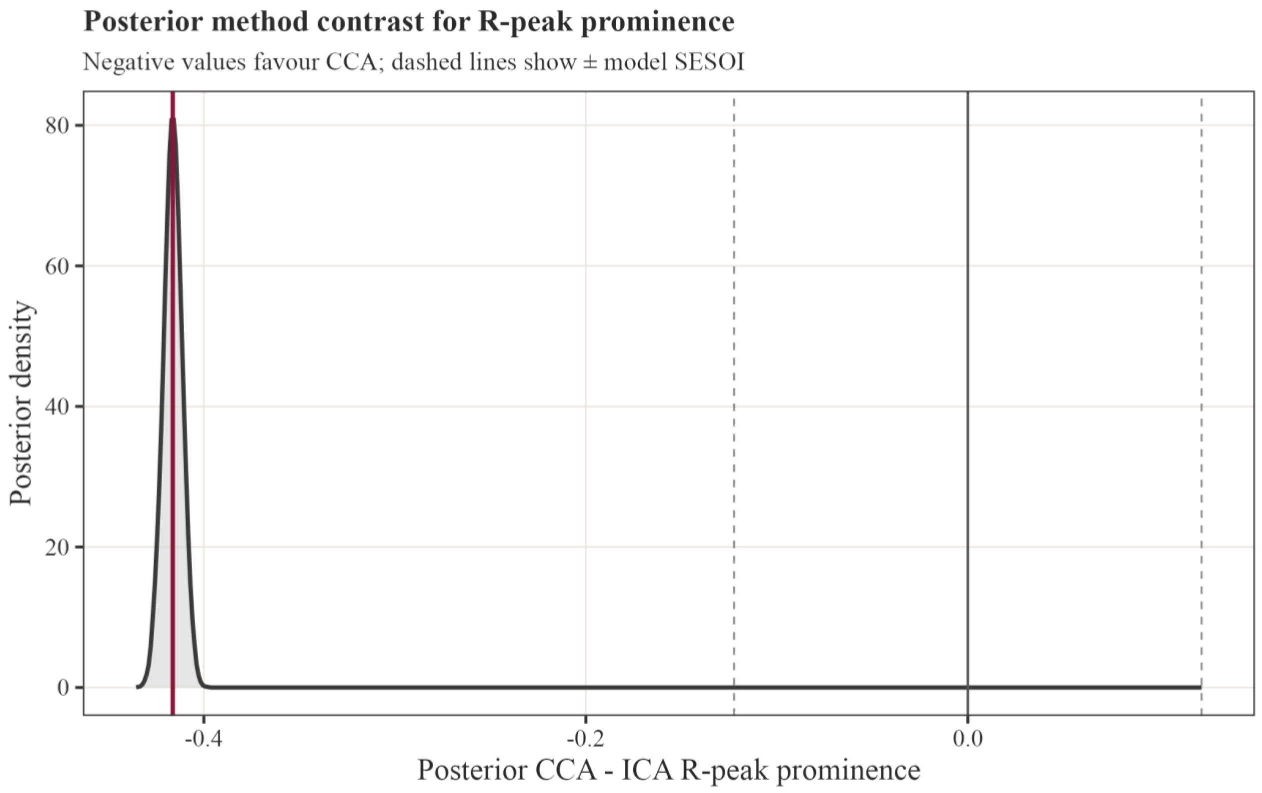


*Figure S7.* Posterior CCA-ICA contrast for alpha power.

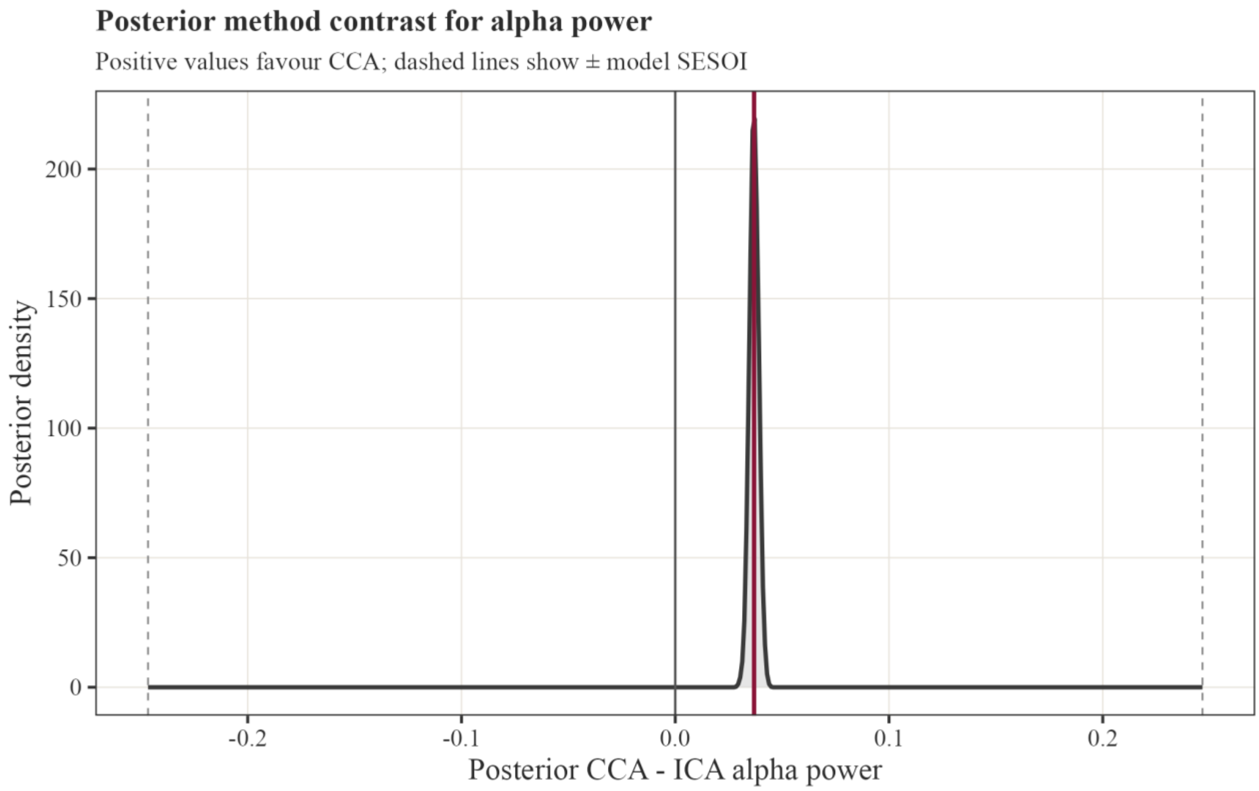


### 5. Stopping point analysis

*Table S6.* Overall stopping-point summaries for ICA and CCA.

| Method | 1st component useful | Max useful | First non-useful stop | All meaningful candidates | Last meaningful stop | Model result |
|---|---|---|---|---|---|---|
| ICA | 56.3% | 56.3% at comp. 1 | 1 [0–1], mode 0 | 3 [2–5], mode 3 | 11 [8–13], mode 11 | 1 useful component; plateau from comp. 2 |
| CCA | 61.3% | 68.3% at comp. 3 | 1 [0–2], mode 0 | 4 [3–5], mode 4 | 9 [4–13], mode 4 | 4 useful components; plateau from comp. 5 |

*Table S7.* Observed stopping transitions by removed component.

| Comp. removed | ICA useful/same/harm % | ICA med Δ SESOI | CCA useful/same/harm % | CCA med Δ SESOI |
|---|---|---|---|---|
| 1 | 56.3/26.5/17.2 | -2.45 | 61.3/19.3/19.4 | -2.31 |
| 2 | 31.0/47.0/22.0 | -0.08 | 56.5/25.5/18.0 | -1.75 |
| 3 | 28.0/58.2/13.8 | -0.15 | 68.3/18.7/13.0 | -3.52 |
| 4 | 27.6/62.8/9.6 | -0.08 | 55.1/34.8/10.1 | -1.38 |
| 5 | 19.7/65.8/14.5 | 0.00 | 20.0/70.3/9.7 | -0.05 |
| 6 | 20.1/69.9/10.0 | -0.12 | 20.3/72.7/7.0 | -0.04 |

| Comp. removed | ICA useful/same/harm % | ICA med Δ SESOI | CCA useful/same/harm % | CCA med Δ SESOI |
|---|---|---|---|---|
| 7 | 18.6/56.3/25.1 | 0.00 | 16.6/69.3/14.1 | 0.00 |
| 8 | 23.5/59.9/16.6 | -0.08 | 20.9/73.7/5.5 | -0.12 |
| 9 | 13.2/71.7/15.1 | 0.04 | 18.0/78.9/3.1 | -0.10 |
| 10 | 27.3/66.2/6.5 | -0.11 | 13.4/80.8/5.8 | -0.04 |
| 11 | 28.2/53.1/18.7 | -0.12 | 7.2/86.1/6.8 | -0.02 |
| 12 | 16.1/71.7/12.2 | 0.00 | 10.7/81.8/7.5 | 0.02 |
| 13 | 16.9/71.4/11.7 | -0.02 | 14.9/83.1/2.0 | -0.14 |
| 14 | 15.8/76.6/7.6 | -0.03 | 10.7/80.4/8.9 | -0.02 |
| 15 | 7.6/89.7/2.7 | -0.05 | 5.8/91.3/3.0 | -0.03 |

*Table S8.* Model-based stopping transitions by removed component.

| Comp. removed | ICA Δ SESOI [95%] | ICA P useful | ICA model | CCA Δ SESOI [95%] | CCA P useful | CCA model |
|---|---|---|---|---|---|---|
| 1 | -1.90 [-2.41, -1.39] | 0.999 | useful | -2.19 [-2.76, -1.62] | 1.000 | useful |
| 2 | -0.31 [-0.72, 0.09] | 0.000 | plateau | -1.41 [-1.93, -0.91] | 0.943 | useful |
| 3 | -0.27 [-0.68, 0.12] | 0.000 | plateau | -2.53 [-2.98, -2.10] | 1.000 | useful |
| 4 | -0.53 [-0.91, -0.15] | 0.007 | plateau | -2.23 [-2.58, -1.89] | 1.000 | useful |
| 5 | -0.12 [-0.49, 0.25] | 0.000 | plateau | -0.38 [-0.69, -0.04] | 0.000 | plateau |
| 6 | -0.17 [-0.53, 0.21] | 0.000 | plateau | -0.26 [-0.59, 0.07] | 0.000 | plateau |
| 7 | 0.09 [-0.26, 0.45] | 0.000 | plateau | -0.18 [-0.50, 0.14] | 0.000 | plateau |
| 8 | -0.24 [-0.59, 0.12] | 0.000 | plateau | -0.49 [-0.80, -0.18] | 0.001 | plateau |
| 9 | 0.12 [-0.22, 0.47] | 0.000 | plateau | -0.31 [-0.61, -0.01] | 0.000 | plateau |
| 10 | -0.67 [-1.00, -0.32] | 0.026 | plateau | -0.22 [-0.52, 0.08] | 0.000 | plateau |
| 11 | -0.35 [-0.69, -0.00] | 0.000 | plateau | -0.03 [-0.33, 0.26] | 0.000 | plateau |
| 12 | -0.22 [-0.55, 0.12] | 0.000 | plateau | -0.06 [-0.36, 0.25] | 0.000 | plateau |
| 13 | -0.17 [-0.52, 0.16] | 0.000 | plateau | -0.31 [-0.62, 0.00] | 0.000 | plateau |
| 14 | -0.26 [-0.59, 0.07] | 0.000 | plateau | -0.17 [-0.47, 0.14] | 0.000 | plateau |
| 15 | -0.20 [-0.52, 0.13] | 0.000 | plateau | -0.08 [-0.40, 0.23] | 0.000 | plateau |

*Figure S8.* Observed consecutive R-peak changes by removed component.

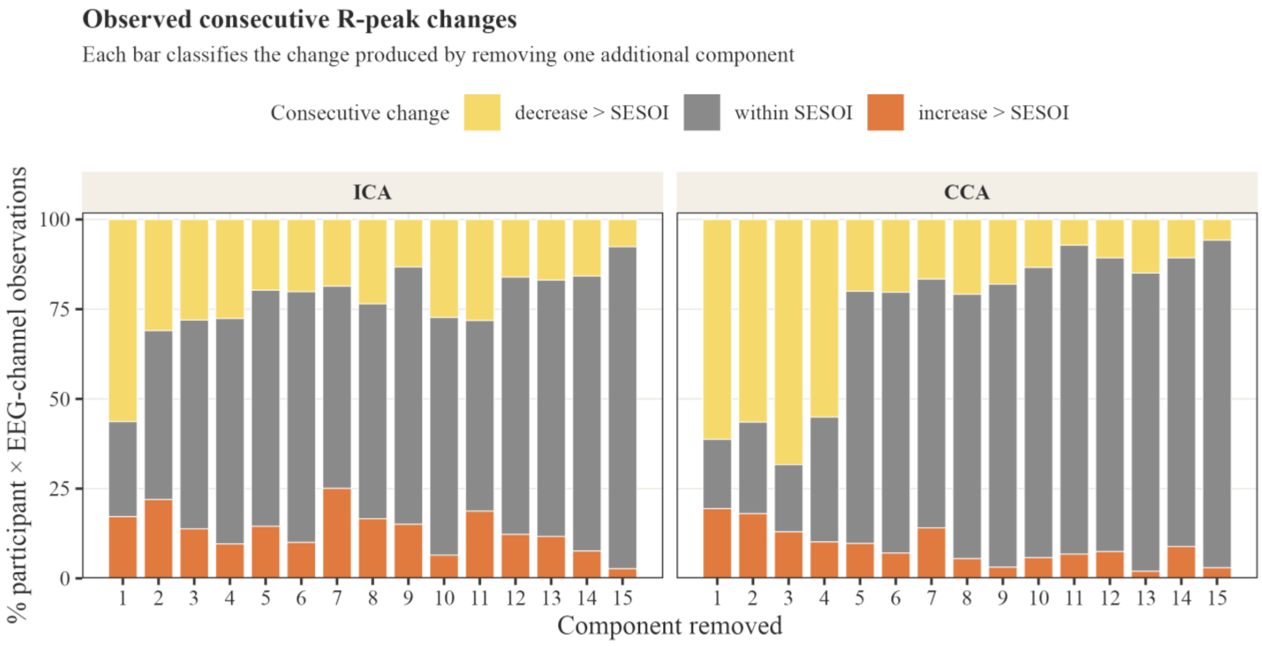


*Figure S9.* Distribution of first non-useful stopping points.

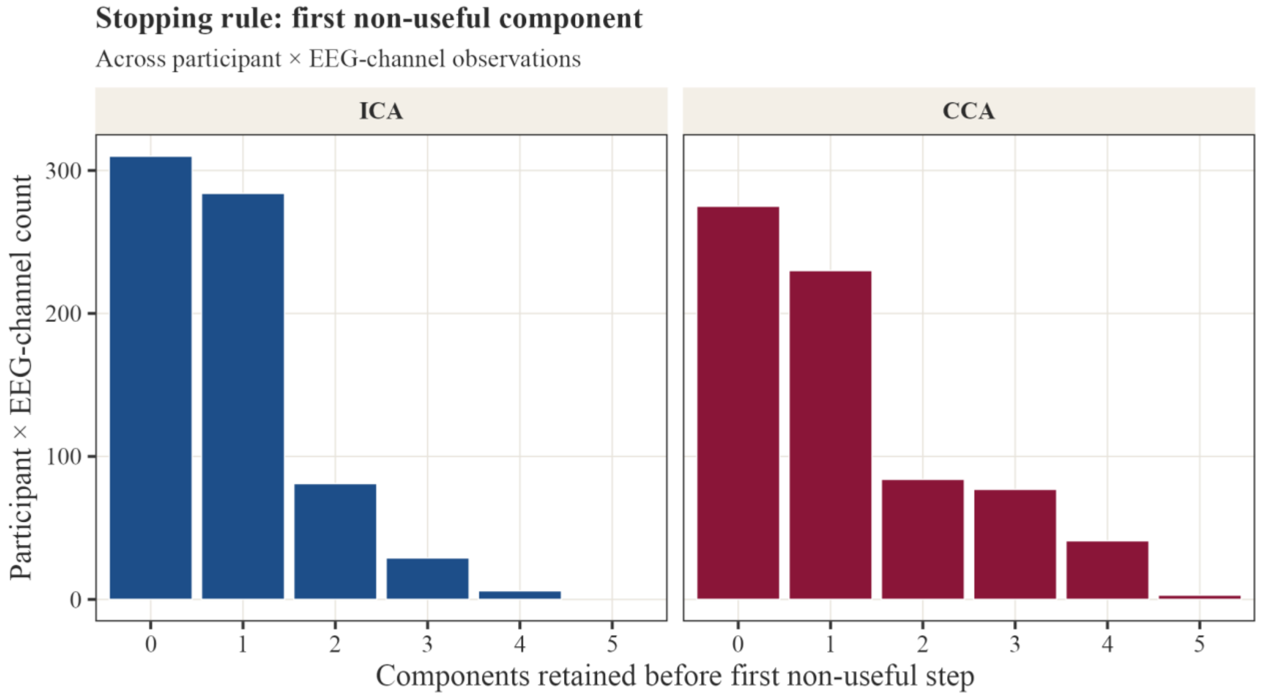


*Figure S10.* Distribution of last meaningful stopping points.

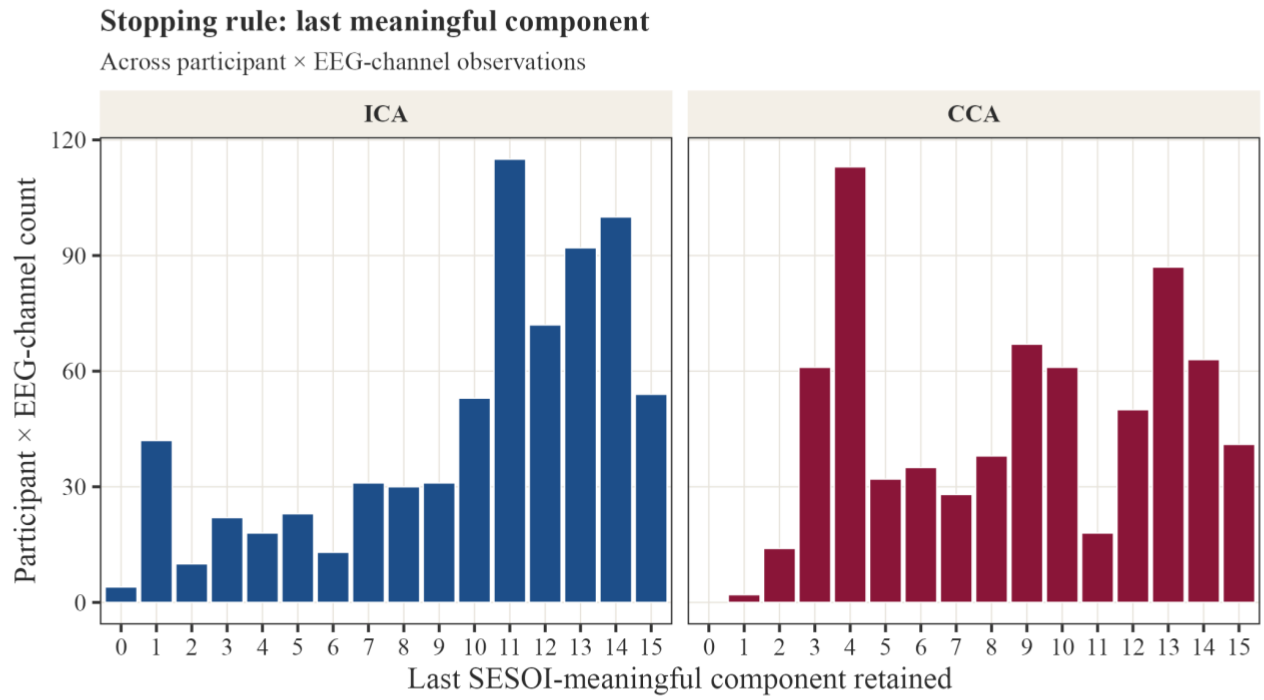


*Figure S11.* Distribution of all SESOI-meaningful component candidates.

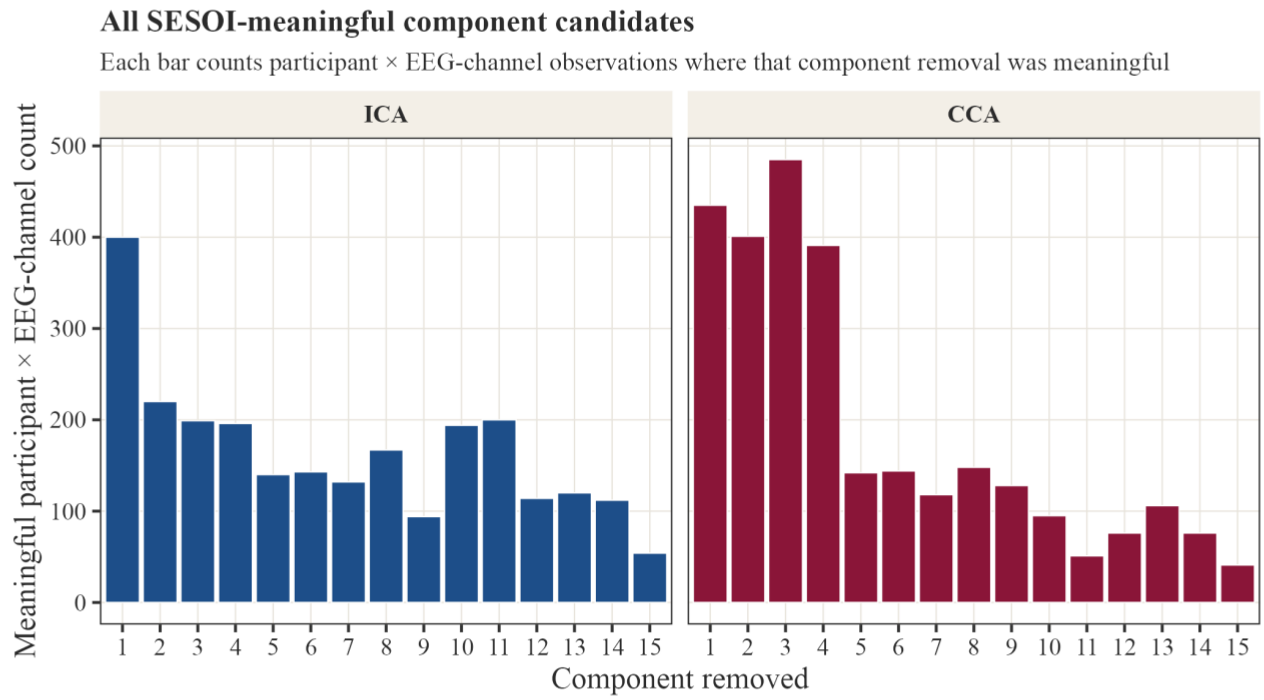


*Figure S12.* Model-estimated consecutive R-peak changes by removed component.

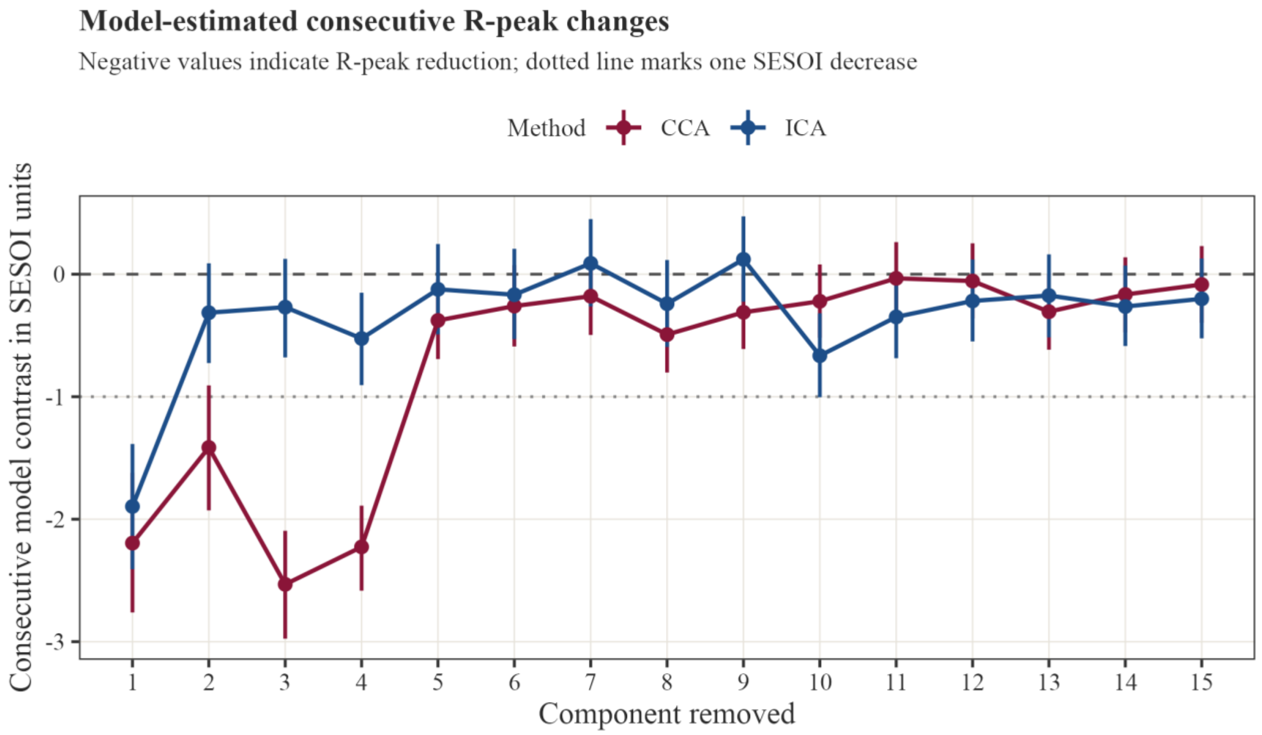


*Figure S13.* Model evidence for useful improvement, plateau, and harm.

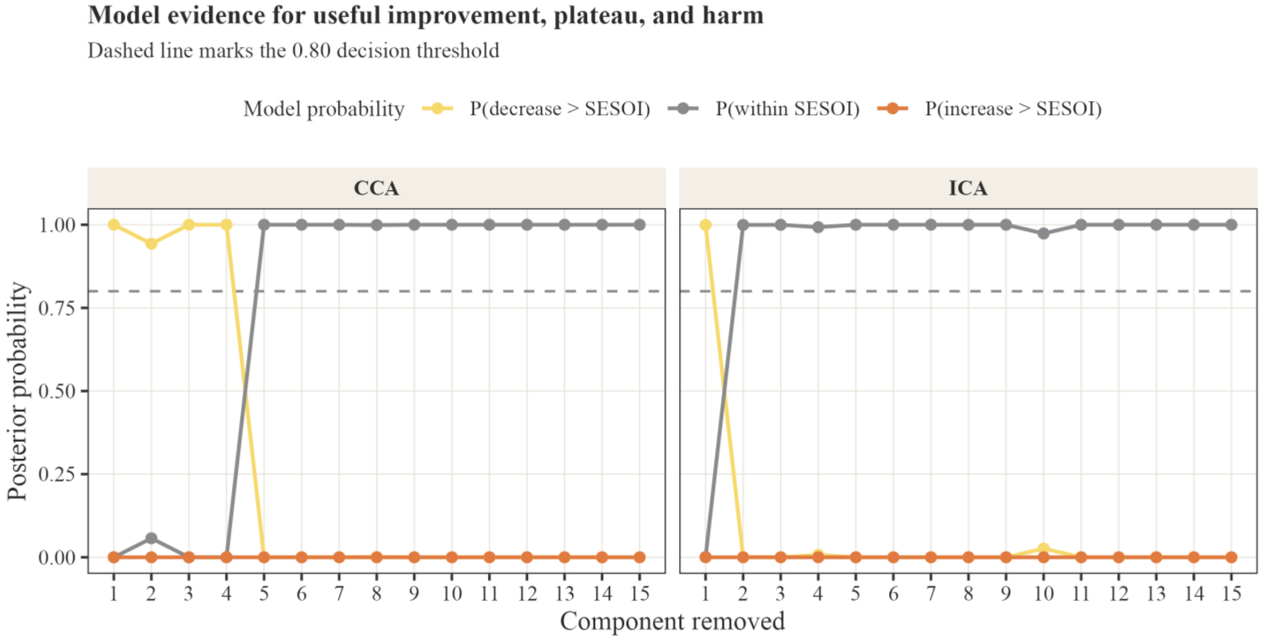


### 6. Top-performing ECG sets for CCA

*Table S9.* Lead-count overview for all exact reduced CCA ECG-channel combinations.

| ECG channels | Exact sets | As well as full15 | Worse than full15 | Loss med/q90, SESOI | Loss med, baseline % | Full15 recovery |
|---|---|---|---|---|---|---|
| 1 | 15 | 3.0% | 97.0% | 10.22 / 16.71 | 51.12% | 32.08% |
| 2 | 105 | 8.2% | 91.8% | 5.80 / 13.58 | 28.99% | 63.52% |

| ECG channels | Exact sets | As well as full15 | Worse than full15 | Loss med/q90, SESOI | Loss med, baseline % | Full15 recovery |
|---|---|---|---|---|---|---|
| 3 | 455 | 16.1% | 83.9% | 3.61 / 11.02 | 18.07% | 77.44% |
| 4 | 1,365 | 25.1% | 74.9% | 2.41 / 8.69 | 12.07% | 85.25% |
| 5 | 3,003 | 34.1% | 65.9% | 1.75 / 6.97 | 8.75% | 89.40% |

*Table S10.* Top three exact CCA ECG-channel sets per channel count.

| n ECG | Rank | Set | Labels | Med ch as-well % | Cells as-well % | P≥50 | Loss med/q90 | Recovery % |
|---|---|---|---|---|---|---|---|---|
| 1 | 1 | 14 | V6 | 3.9 | 4.9 | 0/14 | 8.49/15.00 | 50.0 |
| 1 | 2 | 4 | L supra | 3.0 | 5.1 | 0/14 | 7.49/15.15 | 52.6 |
| 1 | 3 | 3 | R supra | 3.0 | 4.9 | 0/14 | 8.32/14.86 | 50.7 |
| 2 | 1 | 4,7 | L supra+RA | 12.0 | 12.4 | 0/14 | 5.16/13.35 | 66.4 |
| 2 | 2 | 8,14 | LA+V6 | 10.8 | 10.6 | 0/14 | 5.29/13.87 | 67.8 |
| 2 | 3 | 8,13 | LA+V5 | 10.7 | 11.4 | 0/14 | 5.10/12.58 | 70.2 |
| 3 | 1 | 2,3,8 | L front neck+R supra+LA | 26.7 | 23.2 | 0/14 | 2.50/8.13 | 85.0 |
| 3 | 2 | 2,3,4 | L front neck+R supra+L supra | 23.5 | 23.7 | 1/14 | 2.54/8.25 | 84.3 |
| 3 | 3 | 2,7,8 | L front neck+RA+LA | 23.2 | 21.7 | 0/14 | 2.91/8.04 | 82.5 |
| 4 | 1 | 8,10,13,14 | LA+V2+V5+V6 | 40.4 | 37.0 | 2/14 | 1.53/5.14 | 90.5 |
| 4 | 2 | 8,10,12,14 | LA+V2+V4+V6 | 40.0 | 36.5 | 3/14 | 1.46/4.51 | 90.7 |
| 4 | 3 | 8,11,13,14 | LA+V3+V5+V6 | 38.6 | 33.9 | 3/14 | 1.50/5.14 | 91.3 |
| 5 | 1 | 8,9,10,11,12 | LA+V1+V2+V3+V4 | 55.8 | 49.6 | 8/14 | 0.85/3.51 | 95.3 |
| 5 | 2 | 10,11,12,14,15 | V2+V3+V4+V6+LL | 50.0 | 47.2 | 7/14 | 1.01/3.87 | 93.6 |
| 5 | 3 | 3,9,10,11,12 | R supra+V1+V2+V3+V4 | 50.0 | 45.1 | 7/14 | 1.03/4.24 | 93.8 |

*Figure S14.* Reduced-channel CCA performance by ECG-channel count.

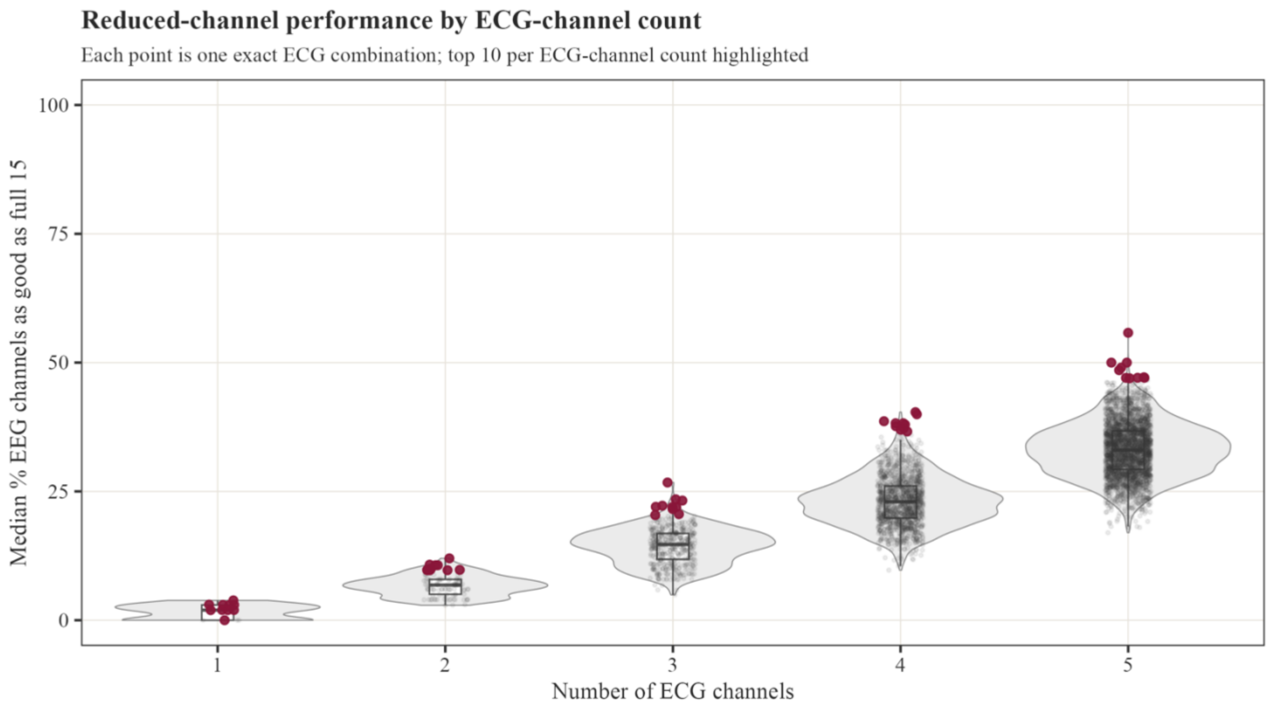


*Figure S15.* Loss of the best CCA candidate by ECG-channel count.

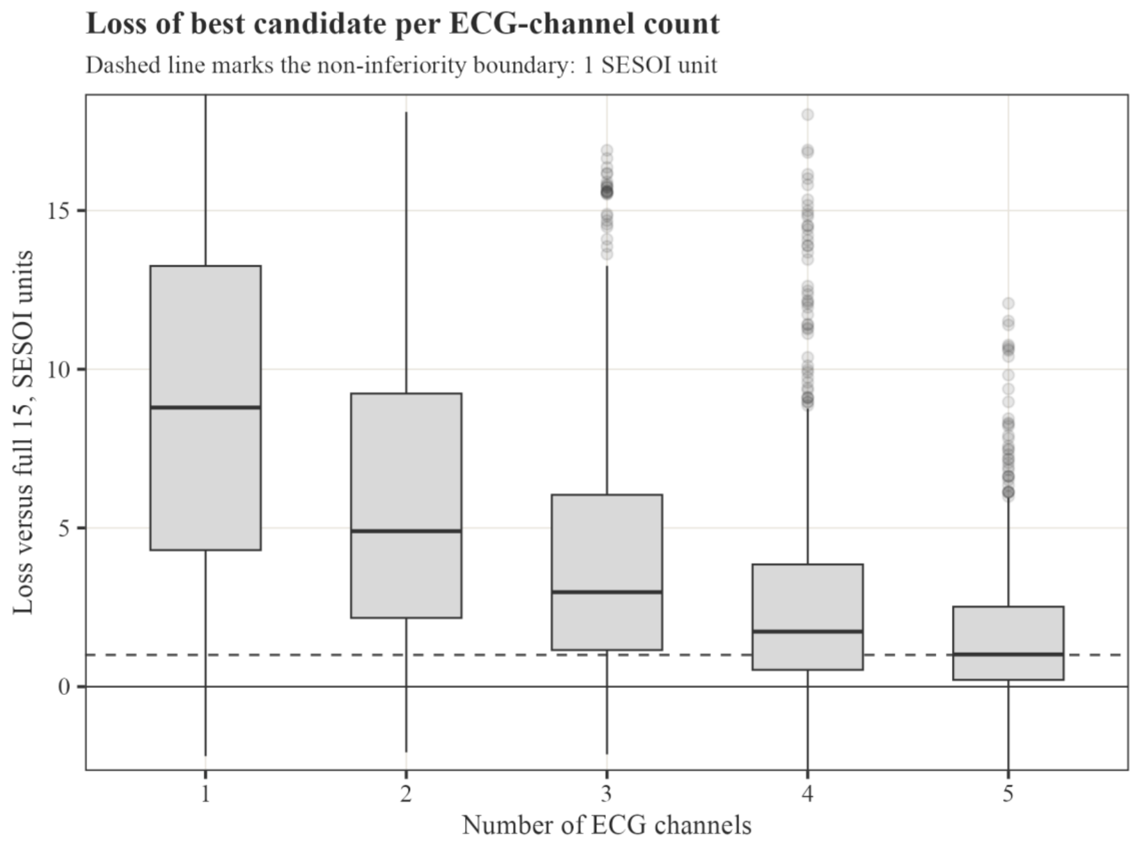


*Figure S16.* Recovery of full 15-channel CCA improvement for the best candidates.

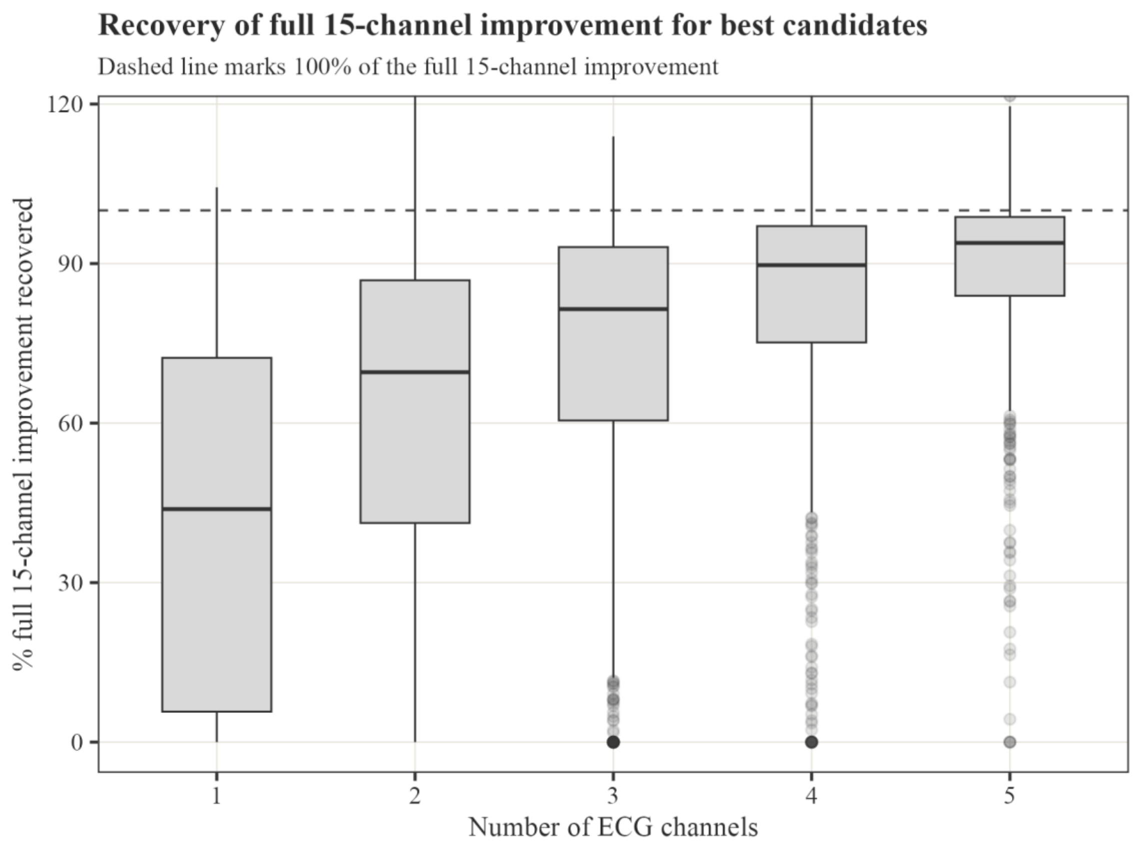


*Figure S17.* Participant robustness of top reduced CCA ECG combinations.

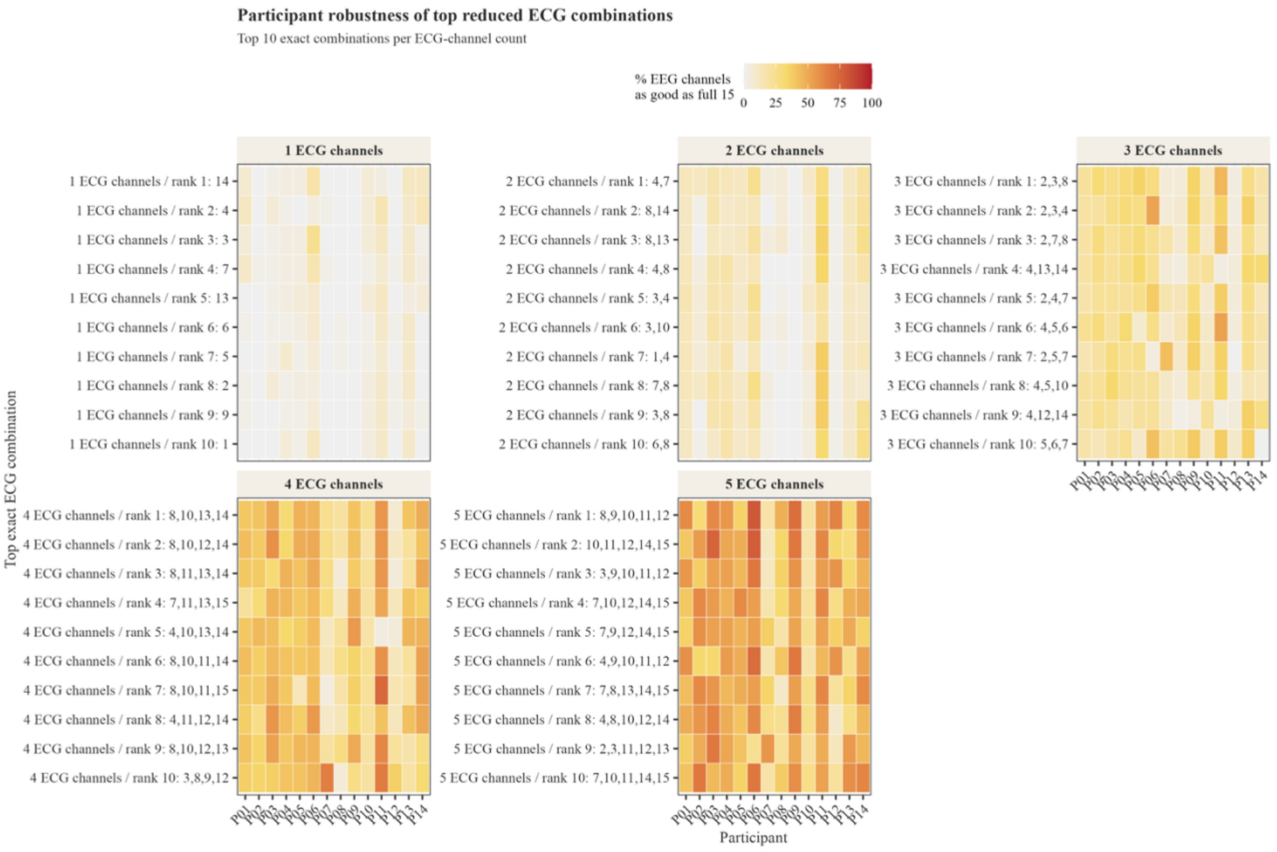


*Figure S18.* EEG-channel robustness of top reduced CCA ECG combinations.

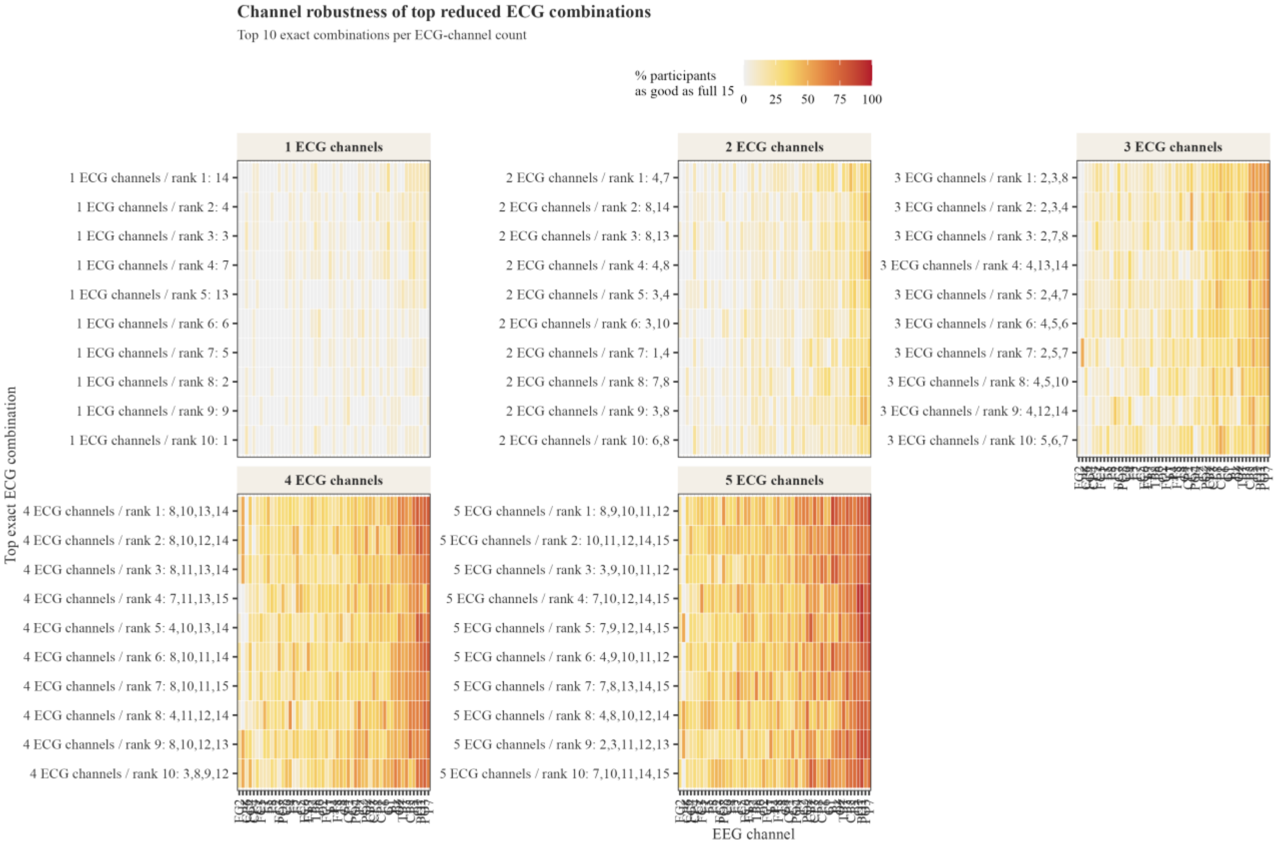


*Figure S19.* Participant-by-EEG-channel robustness for top five-channel CCA candidates.

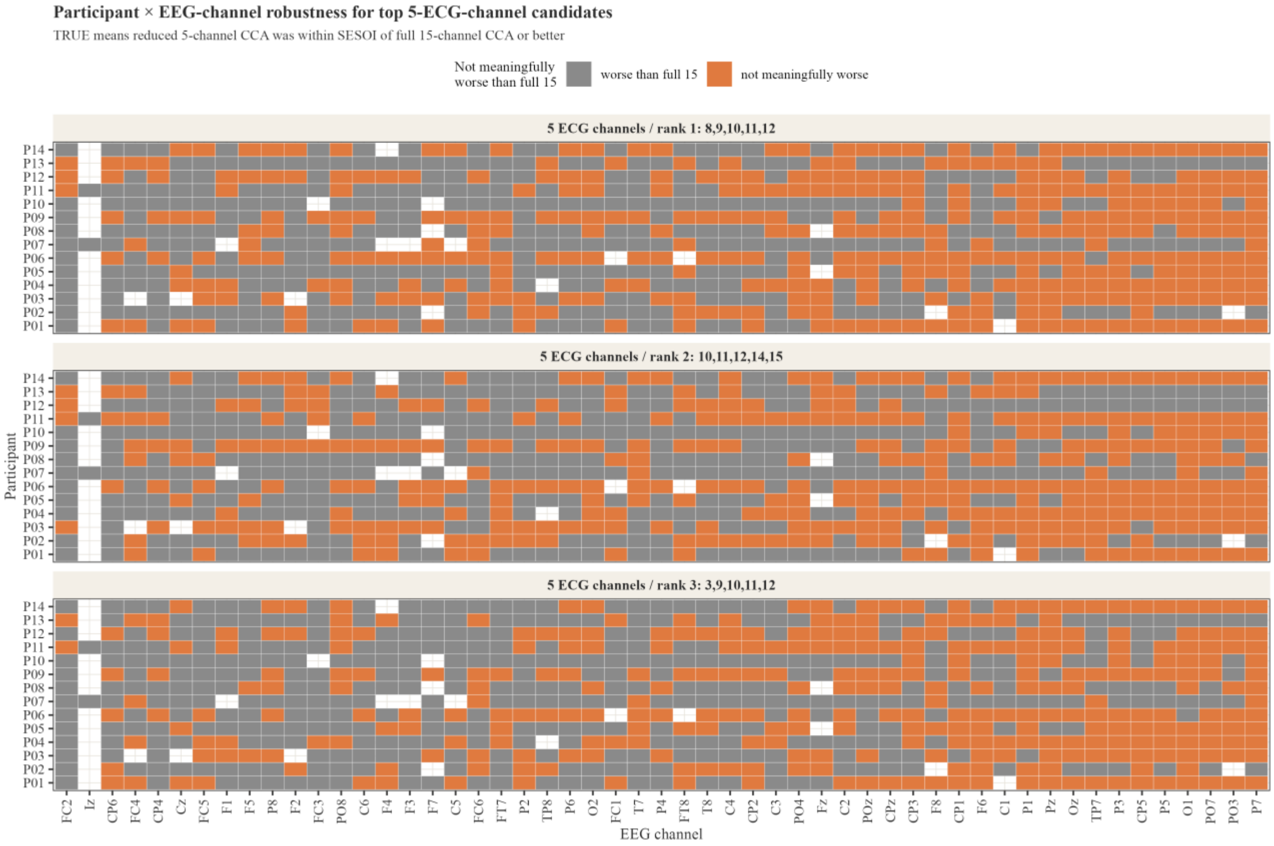


*Figure S20.* Top-20% recurrence of original exact CCA candidates after resampling.

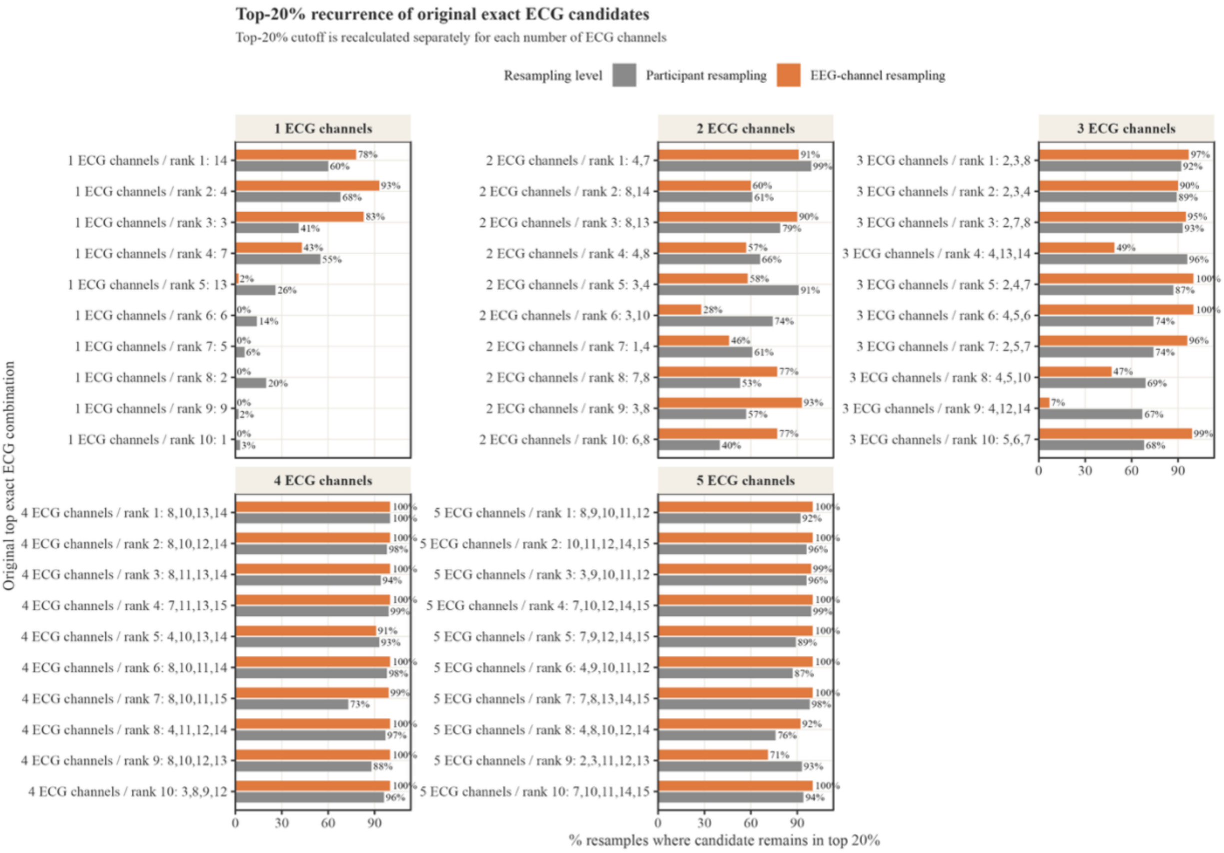


*Figure S21.* Recurring exact subsets among top reduced CCA combinations.

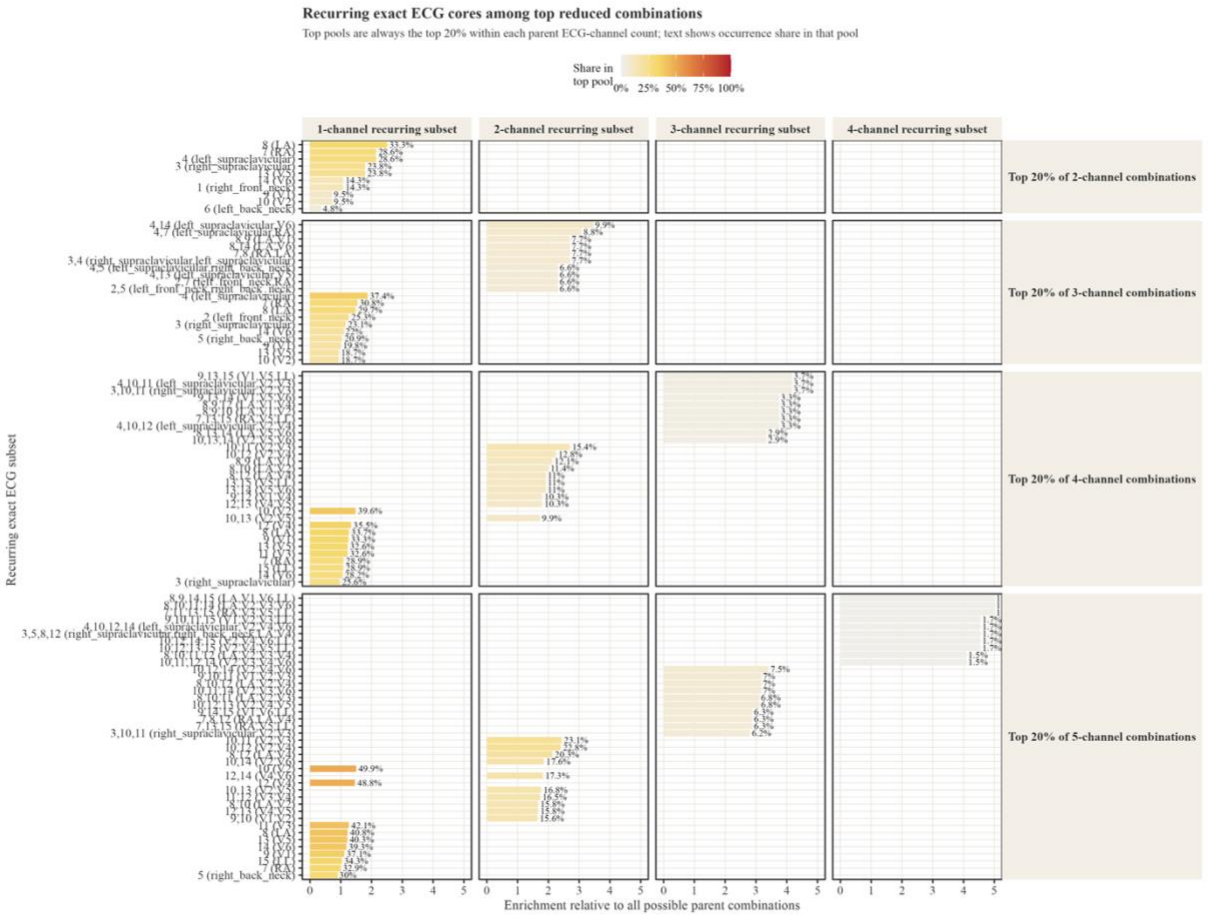


*Figure S22.* Exclusive anatomical-family prevalence among high-ranking CCA combinations.

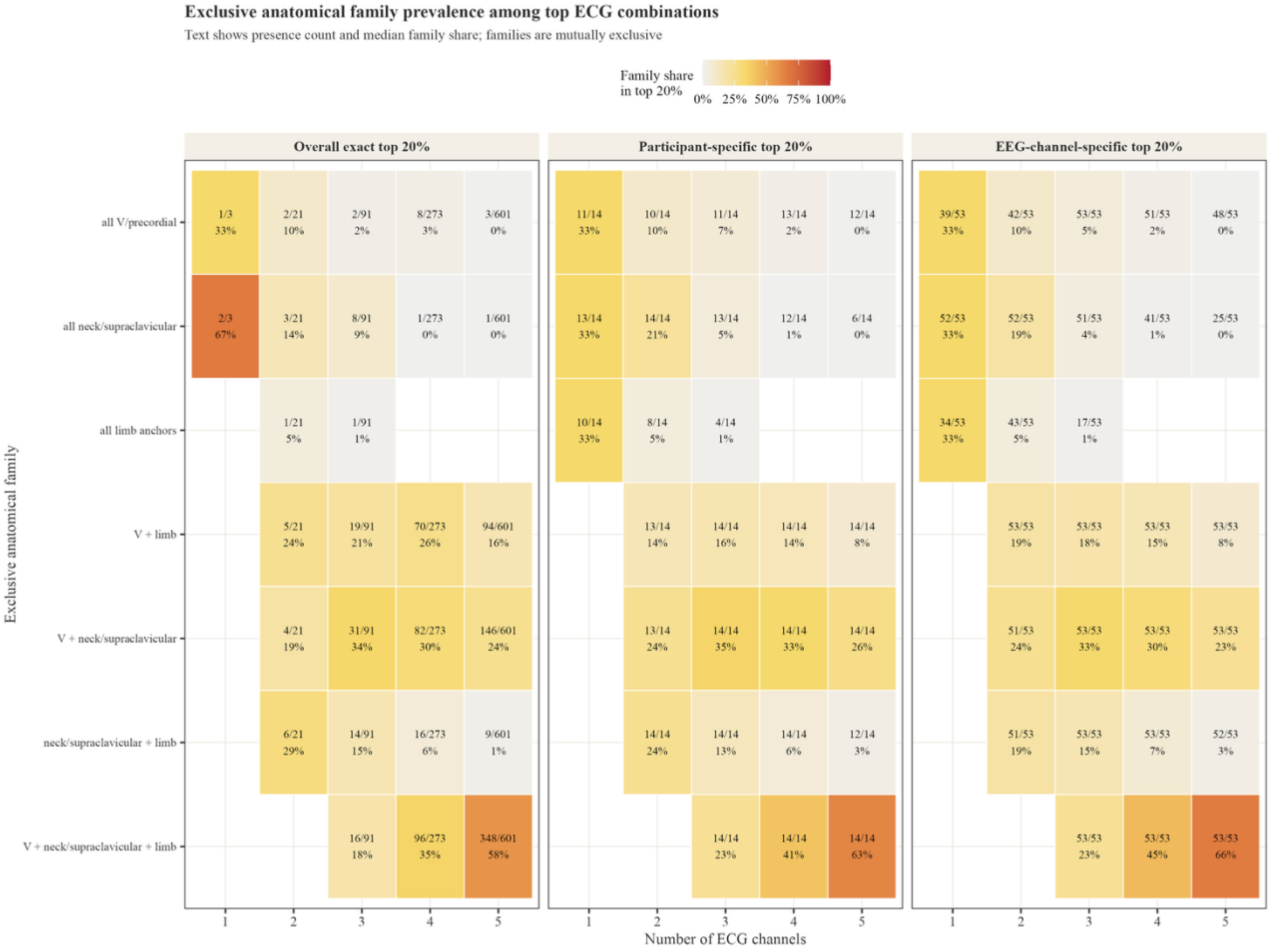


*Figure S23.* Bootstrap anatomical-family stability among top CCA combinations.

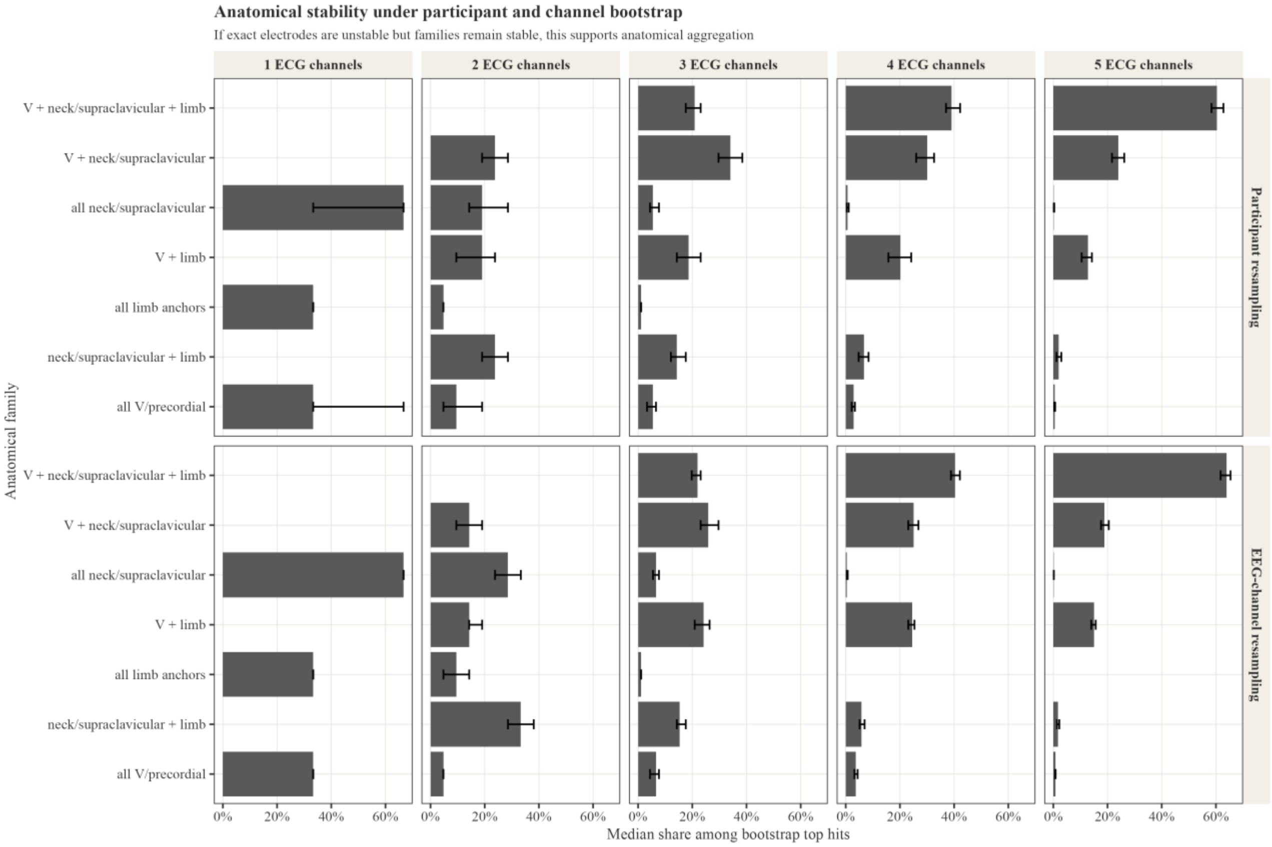

### 7. Top-performing ECG sets for ICA

*Table S11.* Best reduced ICA ECG-channel sets per channel count, scored against the full 15-channel CCA benchmark.

| ECG channels | Best ICA set | Anatomical labels | Median coverage | Participants ≥50% channels | Median loss, SESOI | Full15 CCA recovery |
|---|---|---|---|---|---|---|
| 1 | 9 | V1 | 0.0% | 0/14 | 11.44 | 19.7% |
| 2 | 7,9 | RA + V1 | 2.0% | 0/14 | 8.99 | 35.2% |
| 3 | 10,11,13 | V2 + V3 + V5 | 3.9% | 0/14 | 9.17 | 38.1% |
| 4 | 3,7,8,12 | right supraclavicular + RA + LA + V4 | 4.0% | 0/14 | 6.98 | 51.7% |
| 5 | 9,11,12,14,15 | V1 + V3 + V4 + V6 + LL | 5.8% | 0/14 | 8.28 | 49.7% |

*Figure S24.* Top exact ICA reduced ECG-channel combinations against the full 15-channel CCA benchmark.

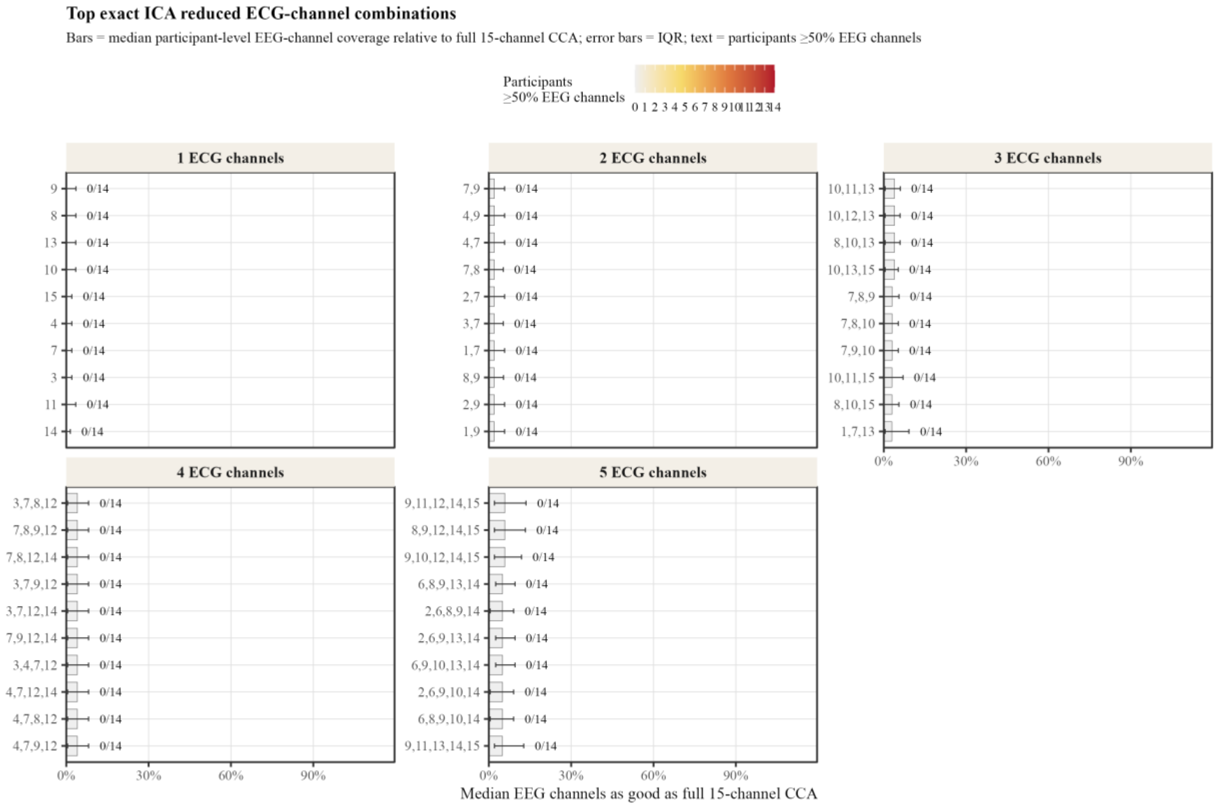


*Figure S25.* ICA reduced-channel performance by ECG-channel count across all exact combinations.

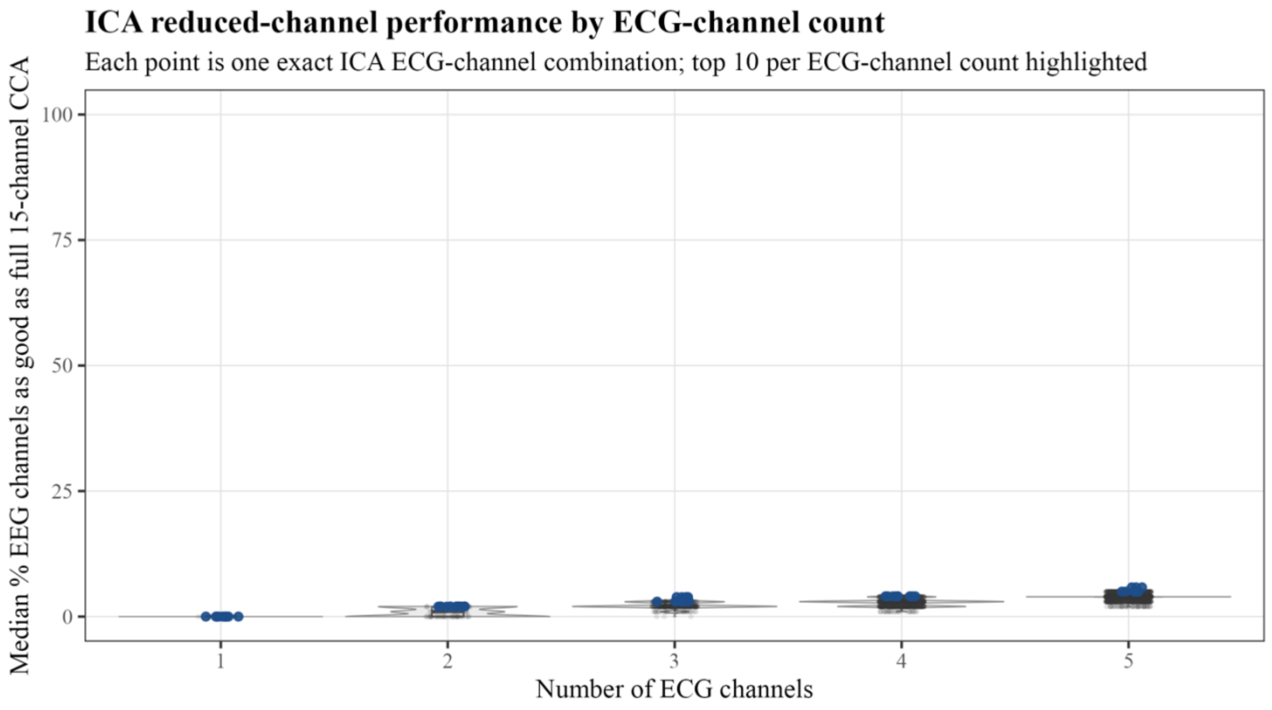


*Figure S26.* Loss of the best ICA candidate by ECG-channel count relative to full 15-channel CCA.

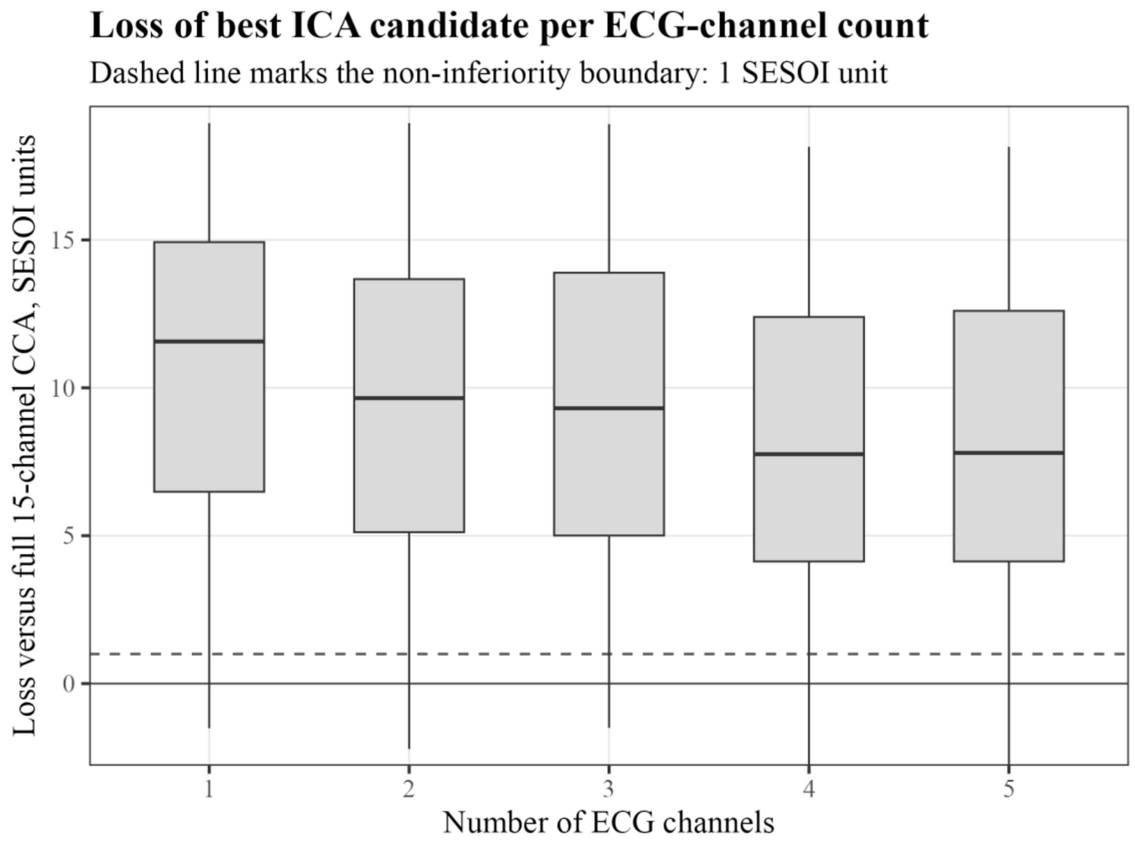


*Figure S27.* Recovery of full 15-channel CCA improvement by reduced ICA ECG-channel count.

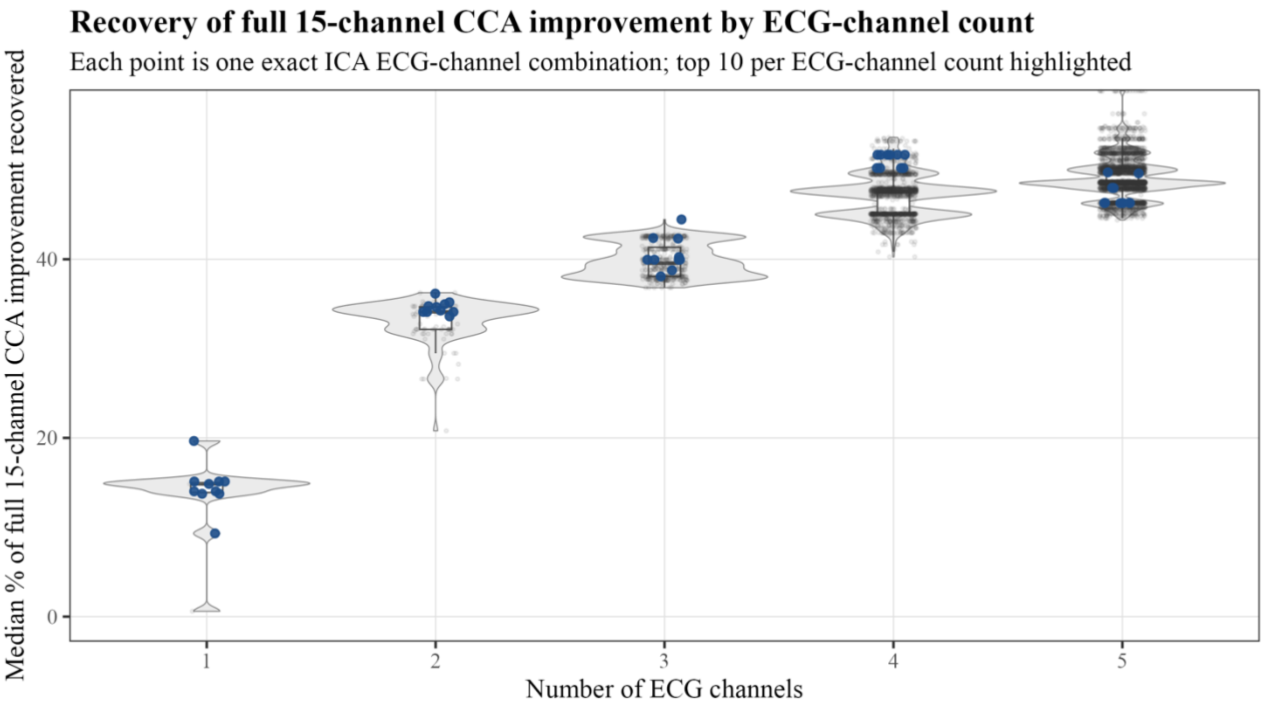


*Figure S28.* Participant-by-EEG-channel robustness of the top five-channel ICA candidates.

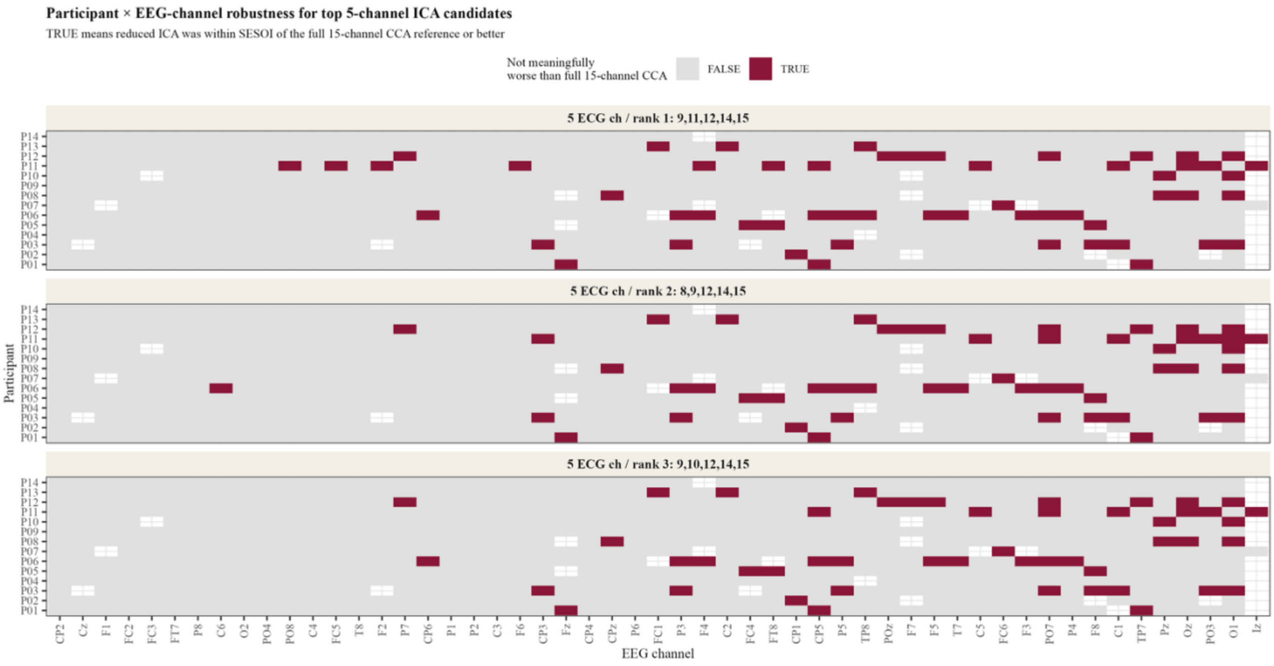


*Figure S29.* Exclusive anatomical-family prevalence among top ICA combinations.

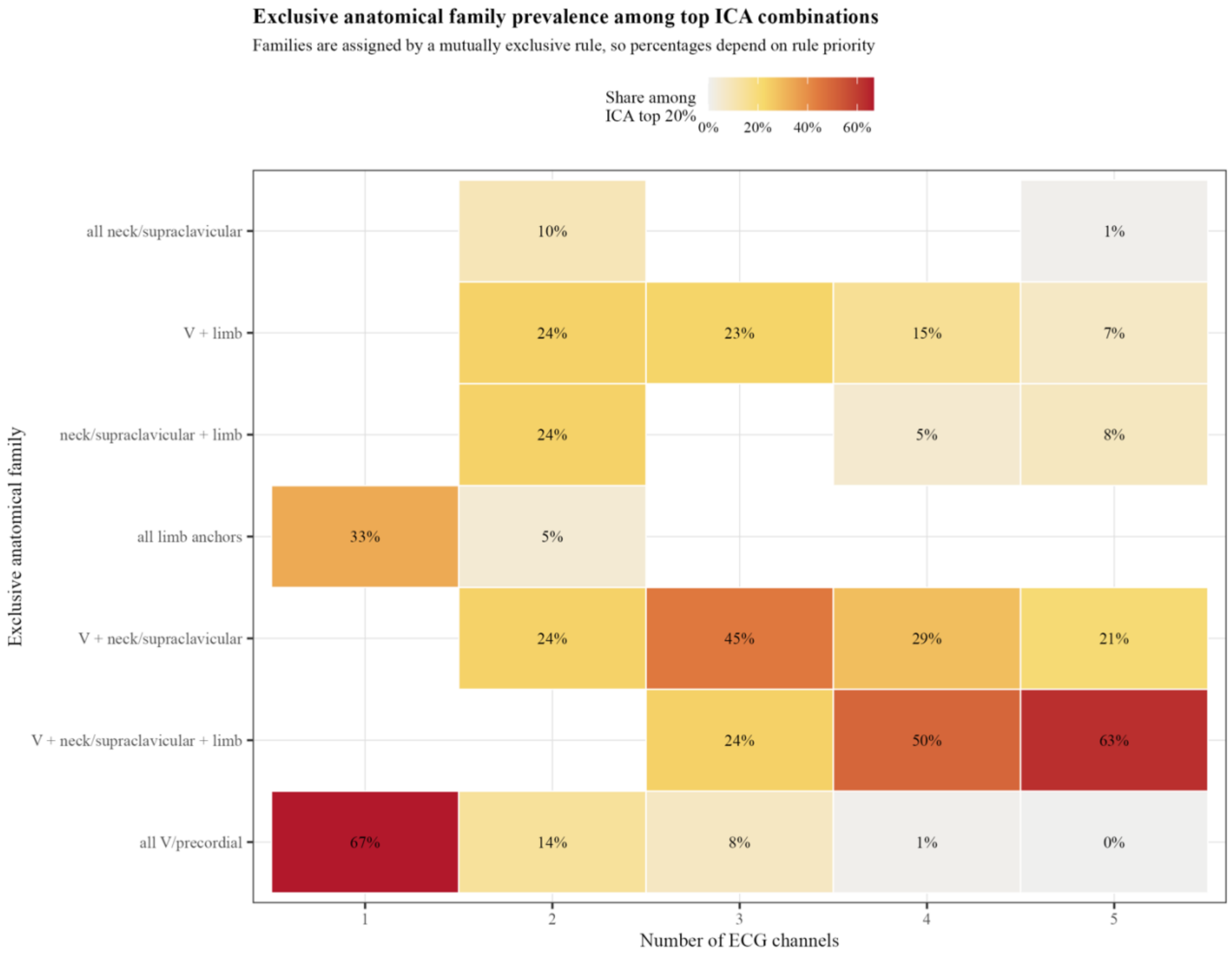


*Figure S30.* Best ICA versus best CCA reduced ECG-channel candidates by channel count.

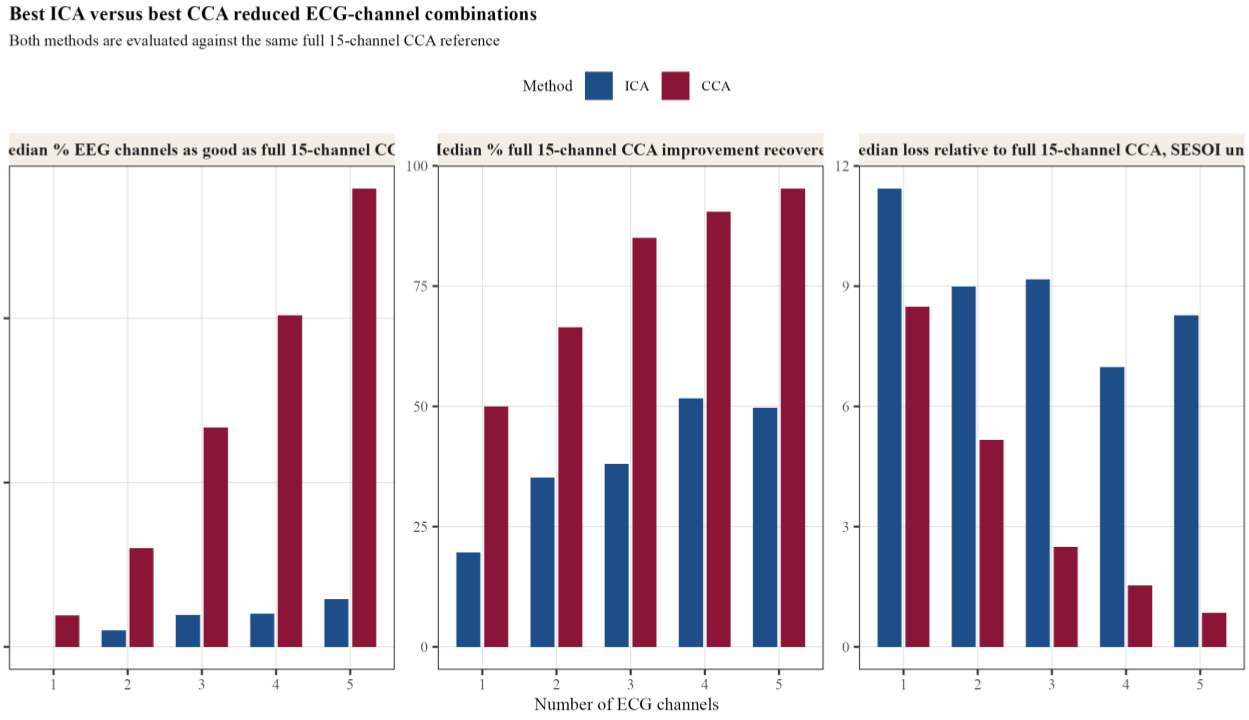


*Figure S31.* Distribution of exact-combination performance under ICA and CCA.

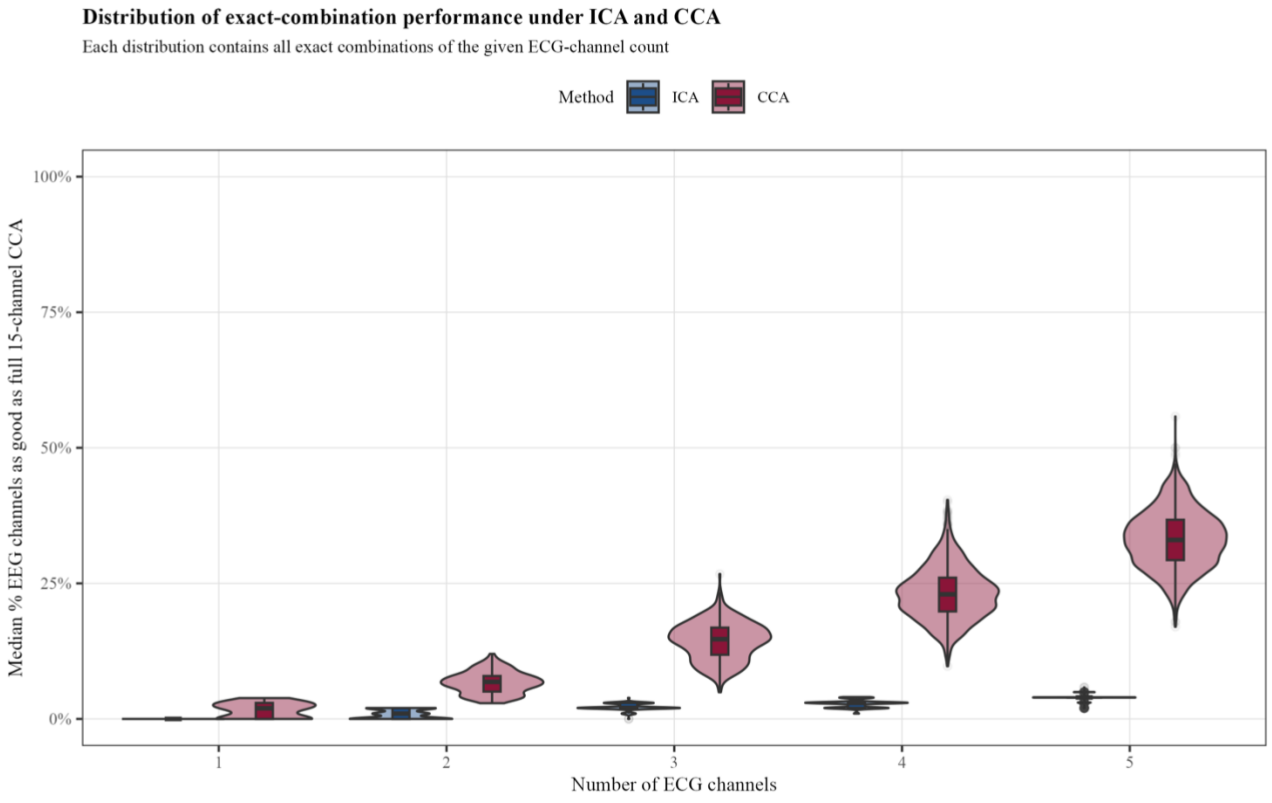


*Figure S32.* Performance of the top CCA-selected ECG combinations when evaluated under ICA.

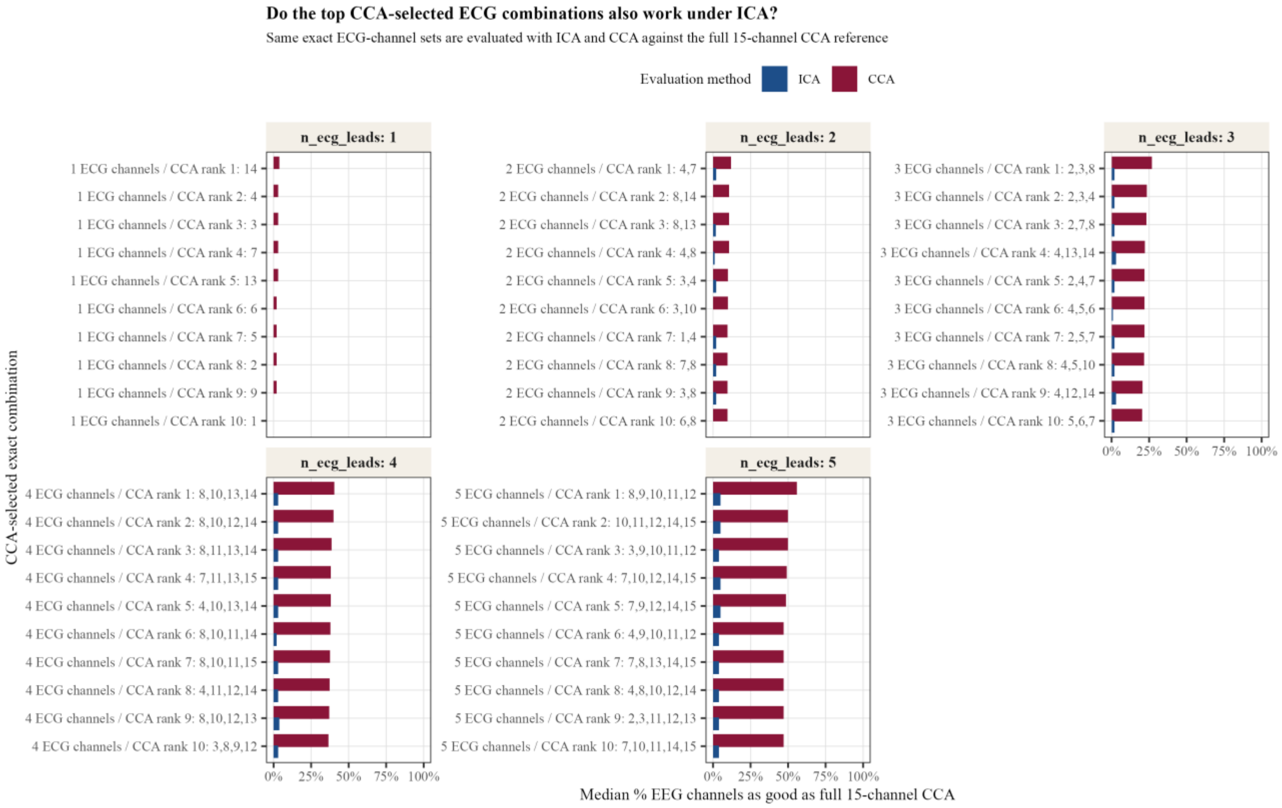
